# PtdIns(3,5)P_2_ Is an Endogenous Ligand of STING in Innate Immune Signaling

**DOI:** 10.64898/2025.12.29.696917

**Authors:** Jay Xiaojun Tan, Bo Lv, Jie Li, Tuo Li, Fenghe Du, Xiang Chen, Xuewu Zhang, Xiao-chen Bai, Zhijian J. Chen

## Abstract

Cytosolic DNA exposure triggers innate immune responses through cyclic GMP-AMP (cGAMP) synthase (cGAS)^1–3^. Upon binding to DNA, cGAS is activated to produce cGAMP, which functions as a second messenger that binds to stimulator of interferon genes (STING), an endoplasmic reticulum (ER)-localized signaling adaptor^3–5^. STING then traffics from the ER to the Golgi, leading to activation of the kinases TBK1 and IKK and subsequent induction of interferons and other cytokines^6–10^. Here we show that phosphatidylinositol 3,5-bisphosphate [PtdIns(3,5)P_2_] is an endogenous ligand of STING that functions together with cGAMP to induce STING activation. Proteomics analysis identified a constitutive interaction between STING and PIKfyve, an enzyme that produces PtdIns(3,5)P_2_ in mammalian cells. Deletion of PIKfyve blocked STING trafficking from the ER and TBK1 activation. *In vitro* reconstitution revealed a strong and selective effect of PtdIns(3,5)P_2_ on STING activation. Purified STING bound directly to PtdIns(3,5)P_2_ in a fluorescence resonance energy transfer (FRET) assay. Consistently, PtdIns(3,5)P_2_ promoted cGAMP-induced STING oligomerization by binding to a groove between STING dimers as revealed by cryo-EM (Li et al., co-submitted). Similar to PIKfyve depletion, mutation of the PtdIns(3,5)P_2_-binding residues in STING blocked its trafficking and downstream signaling. These results reveal PtdIns(3,5)P_2_ as a lipid ligand of STING with essential roles in innate immunity.

## Main text

Cytosolic exposure of DNA is a danger signal that triggers inflammation in microbial infections, autoimmune disease, aging, and age-related degeneration^1^. Cytosolic DNA is detected by cyclic GMP-AMP (cGAMP) synthase (cGAS) which upon DNA binding produces cGAMP as a second messenger^2,3^. cGAMP directly binds to and activates the endoplasmic reticulum (ER)-anchored signaling adaptor stimulator of interferon genes (STING)^3–5^. Upon cGAMP binding, STING traffics from the ER through the Golgi complex to post-Golgi vesicles, accompanied by the activation of TANK-binding kinase 1 (TBK1), a serine/threonine-protein kinase^6–9^. TBK1 triggers the nuclear translocation of both interferon regulatory transcription factor 3 (IRF3) and Nuclear factor kappa-light-chain-enhancer of activated B cells (NF-κB), leading to the transcriptional upregulation of type I interferons and inflammatory cytokines^6–9,11–13^. Both the ER exit of STING and the subsequent TBK1 activation require the oligomerization of STING induced by cGAMP binding^14–16^. It is unclear why TBK1 activation requires STING trafficking from the ER to perinuclear vesicle clusters.

To dissect how STING trafficking activates TBK1, we examined the dynamics of STING trafficking in human BJ-5ta fibroblasts, an established cell model for the study of STING signaling^7^. Digitonin-mediated cGAMP delivery^17^ triggered robust STING trafficking within 20 min (**Extended Data Fig. 1a**). TBK1 association with STING was captured by co-immunoprecipitation (co-IP) within 10 min after cGAMP delivery, but TBK1 activation as indicated by its phosphorylation at Ser172 (pTBK1-S172) was not obvious until 30 min after stimulation (**Fig. 1a**). Immunofluorescence analysis revealed that, between 20 and 30 min after cGAMP exposure, STING was transported from the cis-Golgi to the trans-Golgi network (TGN), and that by 60 min most STING accumulated at perinuclear vesicle clusters in proximity to the Golgi markers (**Fig. 1b**). In immunofluorescence, TGN appeared to be “wrapped” by *cis*-Golgi (**Extended Data Fig. 1b**). As STING traffics through the Golgi complex, it initially overlapped with the cis-Golgi marker GM-130 while “wrapped” around a TGN marker GOLGA4 (**Fig. 1b**, 20’). As STING arrived at the trans-Golgi, it was wrapped by GM-130 but overlapped with GOLGA4 (**Fig. 1b**, 30’). We used GOLGA4 instead of another commonly used TGN marker TGN38 (also known as TGN46) because the latter trafficked out of TGN after cGAMP stimulation^18^ (**Extended Data Fig. 1b, c**). Using an alternative TGN marker, OSBP-PH-GFP, we found a similar pattern of STING trafficking through the Golgi (**Extended Data Fig. 1d**). Consistent with the time-dependent TBK1 phosphorylation observed in immunoblotting (**Fig. 1a**), the staining of phosphorylated TBK1 (p-TBK1-S172) was undetectable at 20 min and detectable at 30 min after cGAMP stimulation (**Fig. 1c**). The pTBK1-S172 signal extensively associated with TGN but was wrapped around by the cis-Golgi (**Fig. 1c**). Similar to STING, p-TBK1 also accumulated at perinuclear vesicles near the Golgi complex by 60 min (**Fig. 1c**). Thus, there is a strong correlation between STING localization to TGN or TGN-derived vesicles and TBK1 phosphorylation (**Fig. 1d**), as previously reported^19–22^. Treatment of BJ cells with Brefeldin A, which is known to block protein trafficking from the ER to Golgi^23^, caused a quick disassembly of TGN (**Extended Data Fig. 1e**). While brefeldin A fully blocked cGAMP-stimulated STING signaling, it did not affect TBK1 interaction with STING in co-IP (**Fig. 1e**), suggesting that STING trafficking is required for TBK1 activation but not TBK1 binding to STING. These results show that STING-mediated TBK1 recruitment and activation are uncoupled, suggesting that STING may require additional factors to activate TBK1 phosphorylation.

**Figure 1.**
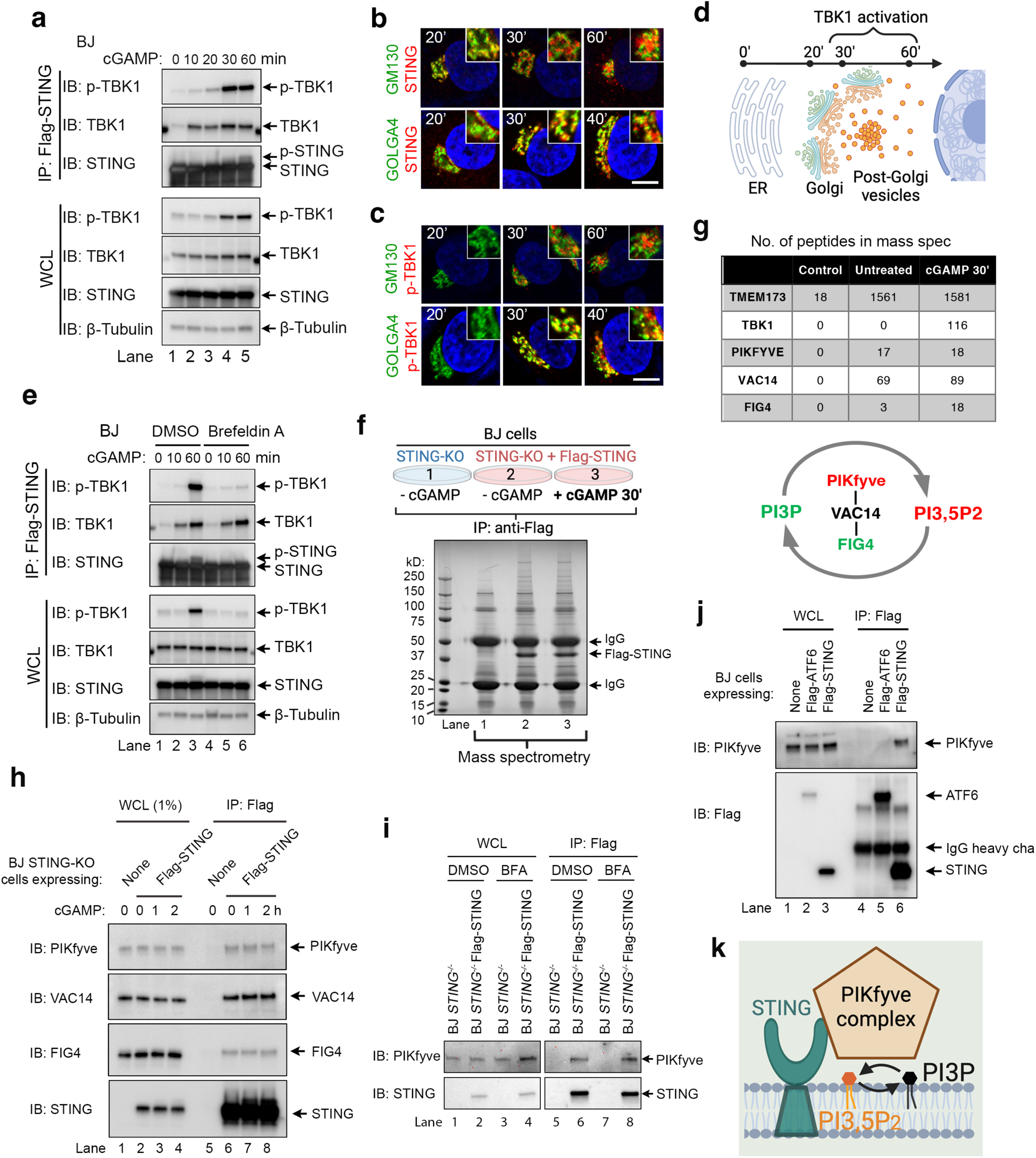
Mass spectrometry identifies STING association with the PIKfyve complex. **(a)** cGAMP stimulates TBK1 phosphorylation and the binding between STING and TBK1. BJ Cells stably expressing Flag-STING were stimulated with 100 nM of cGAMP and whole cell lysates (WCL) were harvested at various time points for immunoprecipitation (IP) with anti-Flag M2 affinity gel. Lysates and IP samples were analyzed by Western blot. **(b, c)** cGAMP stimulates STING trafficking through the Golgi complex. BJ cells were stimulated with 100 nM of cGAMP and fixed at indicated time points for co-staining of (b) STING or (c) phosphor-TBK1 S172 (p-TBK1) with the cis-Golgi Marker GM130 or the TGN marker GOLGA4. Bar, 10 μm. **(d)** Schematic illustration of STING trafficking and TBK1 activation. **(e)** Brefeldin A blocks cGAMP-stimulated TBK1 phosphorylation but not the STING/TBK1 interaction in co-IP. Cells stably expressing Flag-STING were pretreated with 2 μM of Brefeldin A for 1 hour and then stimulated with 100 nM of cGAMP, and whole cell lysates (WCL) were harvested at various time points for immunoprecipitation (IP) with anti-Flag M2 affinity gel. Lysates and IP samples were analyzed by Western blot. **(f)** Mass spectrometry analysis of STING interacting proteins 30 min after cGAMP delivery. The gel image shows silver staining of IP samples used for mass spectrometry analysis. STING knockout (STING-KO) BJ cells stably re-expressing Flag-STING were treated with 1 μM cGAMP and whole cell lysates were harvested 30 min later for IP with anti-Flag M2 affinity gel. STING-KO cells without Flag-STING expression were used as a negative control. **(g)** Top: Peptide numbers of STING and PIKfyve related proteins detected in mass spectrometry. Bottom: illustration of the role of the PIKfyve complex in PtdIns(3,5)P_2_ turnover. **(h)** STING constitutively associates with the PIKfyve complex. BJ STING-KO cells stably expressing Flag-STING were treated with 1 μM cGAMP, and then cell lysates were used for IP with anti-Flag M2 affinity gel, followed by western blot analysis. **(i)** ER-to-Golgi trafficking is not required for the STING-PIKfyve interaction. BJ STING-KO cells stably expressing Flag-tagged STING were pretreated with DMSO or 2 µM Brefeldin A (BFA) for 1 hour to block ER-to-Golgi trafficking. Cell lysates were subjected to IP with anti-Flag M2 affinity gel, followed by western blot analysis. **(j)** PIKfyve selectively associates with STING but not ATF6, another ER-anchored protein. Whole cell lysates of BJ cells stably expressing Flag-tagged ATF6 or STING were subjected to IP with anti-Flag M2 affinity gel, followed by western blot analysis. **(k)** Schematic illustration of STING association with the PIKfyve complex.

### Proteomics identifies the PIKfyve complex as STING-interacting proteins

To search for additional cellular factors that contribute to STING-dependent TBK1 activation, we employed a proteomic approach to analyze the STING interactome after 30 min of cGAMP stimulation (**Fig. 1f**), the minimally required time for cGAMP-induced TBK1 activation (**Fig. 1a**). We affinity-purified stably expressed Flag-tagged STING from human BJ cells and analyzed the associated proteins by mass spectrometry (**Fig. 1f**). As expected, this approach captured cGAMP-dependent enrichment of TBK1 and the Golgi protein ACBD3 in Flag-STING IP samples (**Extended Data Fig. 2a**), consistent with STING interactions with both of them upon cGAMP treatment^24,25^ (**Extended Data Fig. 2b, c**). However, besides TBK1 and ACBD3, no additional proteins were enriched in the STING interactome after cGAMP treatment (**Extended Data Fig. 2a**). We thus switched our focus to proteins constitutively associated with STING.

One major difference among subcellular organelles is the uneven distribution of different membrane lipids; compared with the ER and the cis-Golgi, peripheral compartments including TGN, endolysosomes, and the plasma membrane are more enriched in cholesterol and negatively charged phospholipids such as phosphatidylserine (PS) and phosphoinositides^26,27^. These charged lipids are minor components of cellular membranes but they orchestrate various cellular processes, such as membrane trafficking, cell signaling, and gene expression^28^. We explored the proteomics data searching for potential STING association with lipid-related enzymes and noticed constitutive STING association with all three components of the FYVE finger-containing phosphoinositide kinase (PIKfyve) complex (**Fig. 1g**). This complex consists of PIKfyve itself, the Sac1 domain-containing PtdIns(3,5)P_2_ 5-phosphatase Sac3 (FIG4), and the scaffold protein ArPIKfyve (VAC14) (**Fig. 1g**). PIKfyve is the only enzyme known to be responsible for PtdIns(3,5)P_2_ generation in mammalian cells^29–33^, whereas FIG4 coordinates with PIKfyve to control the dynamic turnover of PtdIns(3,5)P ^34,35^. The constitutive association of STING and the PIKfyve complex was confirmed in co-IP assays, which appeared to be a relatively weak association as only 1% of total PIKfyve was co-precipitated with STING (**Fig. 1h, Extended Data Fig. 2d**). Disrupting the Golgi by Brefeldin A did not affect STING interaction with PIKfyve in co-IP (**Fig. 1i**), indicating that the interaction does not require STING trafficking to the Golgi. This interaction was selective for STING but not another ER-localized transmembrane protein cyclic AMP-dependent transcription factor 6 (ATF6) (**Fig. 1j**). Consistent with constitutive association, the colocalization pattern between STING and PIKfyve did not obviously change before and after STING activation by cGAMP (**Extended Data Fig. 2e**). These results indicate that the PIKfyve complex constitutively and specifically associates with STING, suggesting potential regulation of STING by the lipid messenger PtdIns(3,5)P_2_ (**Fig.1k**).

### PIKfyve is essential for STING signaling

To investigate whether PIKfyve regulates STING signaling, the CRISPR/Cas9 approach was used to deplete PIKfyve in BJ cells. Consistent with a previous report^36^, cells expressing Cas9 and PIKfyve-targeting single guide RNA (sgRNA) developed large lysosomal vacuoles over time (**Extended Data Fig. 3a**). However, most cells with vacuoles were lost in the process of selecting for stable cell clones, suggesting that complete elimination of PIKfyve is lethal. Two single-cell sgRNA clones with mild vacuolation were obtained (**Extended Data Fig. 3b**). Genomic DNA sequencing revealed that these clones carried frame-shift mutations around the sgRNA targeting site in exon 27 of both alleles (**Extended Data Fig. 3c**). However, consistent with the *Pikfyve^β-geo/β-geo^* gene trap mice reported by Zolov et al^37^, we found that the two sgRNA clones that managed to survive were hypomorphs expressing low levels of PIKfyve (**Fig. 2a**), possibly due to alternative splicing around the mutation sites which often occurs in gene trap mice^38,39^. Notably, both clones exhibited increased basal levels of STING, but cGAMP-stimulated phosphorylation of TBK1 and STING appeared normal (**Fig. 2a**). As the remaining amounts of PIKfyve may be sufficient to support STING signaling^33^, small interference RNAs (siRNA) were used to further deplete the remaining PIKfyve protein in these cells. Such near complete depletion of PIKfyve induced the accumulation of more vacuoles (**Extended Data Fig. 3b**) but did not affect cell proliferation within 72 hours (**Extended Data Fig. 3d**). Remarkably, complete loss of PIKfyve blocked STING and TBK1 phosphorylation in response to cGAMP treatment (**Fig. 2b**). Two distinct pairs of siRNAs showed similar efficiency in PIKfyve depletion, and both blocked STING and TBK1 phosphorylation upon cGAMP or herring testis DNA (HT-DNA) treatment (**Fig. 2b, c**). Importantly, PIKfyve siRNAs had effects only in PIKfyve-CRISPR clones but not in wild-type BJ cells (**Fig. 2b, c**), indicating that the effects of these siRNAs on STING signaling were dependent on a complete depletion of PIKfyve. Consistently, PIKfyve siRNAs induced the accumulation of vacuoles only in the PIKfyve sgRNA clones but not in wild-type BJ cells (**Extended Data Fig. 3b**). Notably, complete PIKfyve depletion abolished TBK1 activation selectively in the DNA-sensing pathway (**Fig. 2b, c**) but not the RNA-sensing pathway (Sendai virus, SeV, **Extended Data Fig. 3e**) or the TNFα pathway (**Extended Data Fig. 3f**). Thus, loss of PIKfyve specifically blocked STING-mediated TBK1 signaling without causing a general defect of TBK1 activation (**Extended Data Fig. 3g**).

**Figure 2.**
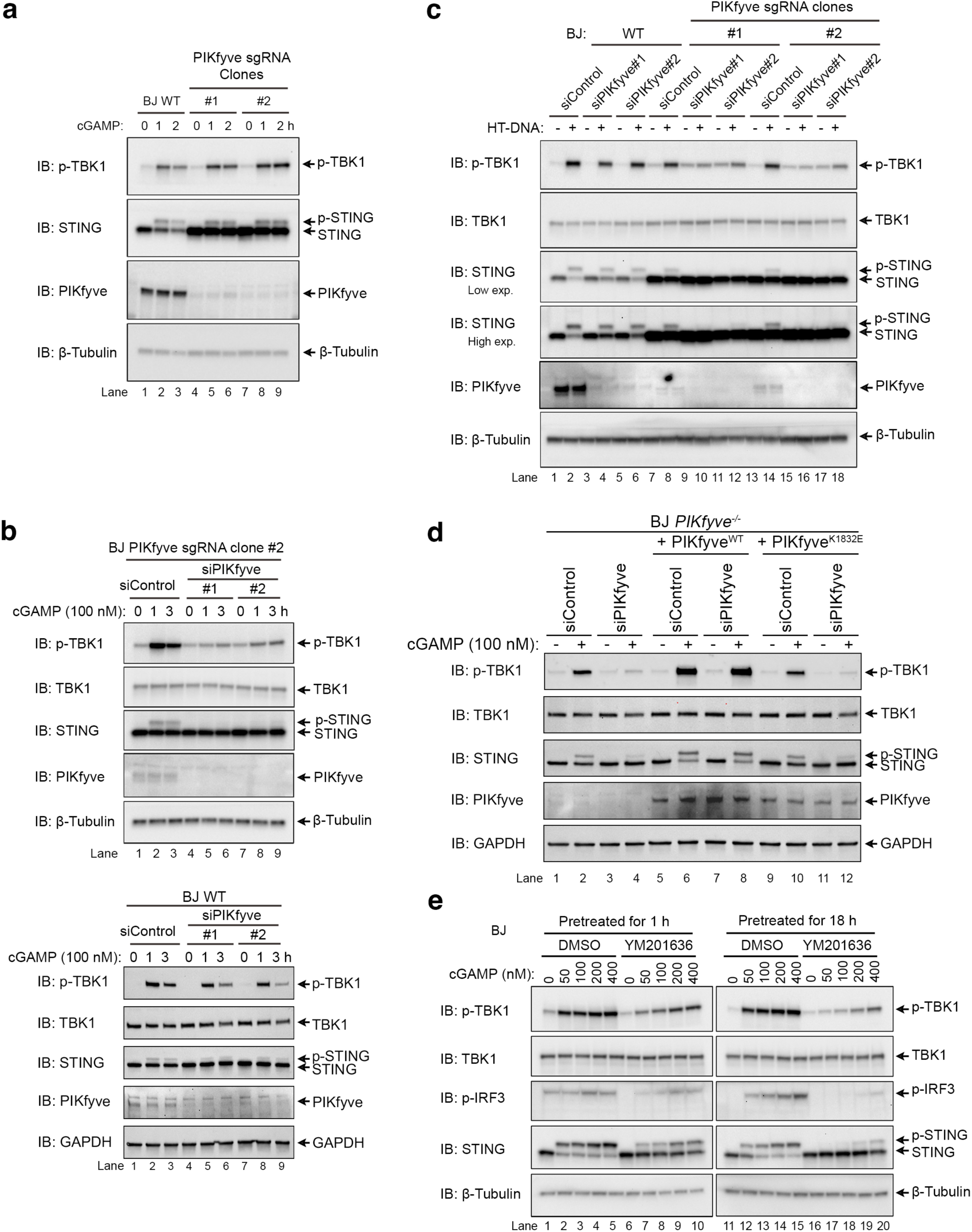
PIKfyve depletion or inhibition blocks STING-mediated TBK1 activation. **(a)** Two BJ PIKfyve sgRNA cell lines with very low levels of PIKfyve have normal STING and TBK1 phosphorylation upon cGAMP treatment. Cells were treated with 100 nM cGAMP and whole cell lysates at indicated time points were analyzed by Western blot. **(b)** RNAi-mediated depletion of remaining PIKfyve in BJ PIKfyve sgRNA cell lines blocks cGAMP-stimulated phosphorylation of STING and TBK1. Cells were transfected with indicated siRNAs; 3 days later cells were treated with 100 nM cGAMP and whole cell lysates harvested at indicated time points were analyzed by Western blot. **(c)** RNAi-mediated depletion of remaining PIKfyve in BJ PIKfyve sgRNA cell lines blocks HT-DNA stimulated phosphorylation of STING and TBK1. Cells were transfected with HT-DNA at a concentration of 2 μg/ml; 3 hours later whole cell lysates were harvested and analyzed by Western blot. **(d)** The defects of STING-TBK1 signaling in PIKfyve-depleted is rescued by the reconstitution of wild-type PIKfyve, but not its kinase dead mutant K1832E. Indicated cells were treated with 100 nM cGAMP for 1 h and whole cell lysates were analyzed by Western blot. **(e)** Prolonged treatment with the PIKfyve inhibitor (YM201636) inhibits cGAMP-stimulated phosphorylation of STING and TBK1 in BJ cells. Cells were pretreated with 5 μM YM201636 for 1 hour or 18 hours before stimulation with indicated concentrations of cGAMP; 1 hour after cGAMP treatment, whole cell lysates were harvested and analyzed by Western blot.

It was previously reported that less than 10% of PIKfyve protein in *Pikfyve^β-geo/β-geo^* gene trap mice was sufficient to generate about 50% of cellular PtdIns(3,5)P_2_, but further depleting the remaining 10% of PIKfyve caused a complete removal of all PtdIns(3,5)P_2_ in *Pikfyve^β-geo/β-geo^* fibroblasts^37^. Since a complete depletion of PIKfyve is necessary to block STING-mediated TBK1 signaling, it is likely that the kinase activity of PIKfyve, i.e., PtdIns(3,5)P_2_ production, plays a role in this pathway. Indeed, reconstitution of wild-type PIKfyve, but not its kinase dead mutant, fully rescued STING-TBK1 signaling in cells depleted of endogenous PIKfyve (**Fig. 2d**). Pretreatment of BJ cells for 1 hour with YM201636, a well characterized small molecule inhibitor of PIKfyve ^40^, partially suppressed cGAMP-stimulated phosphorylation of STING and TBK1 (**Fig. 2e**, left). Prolonged treatment with YM201636, which is known to achieve a better depletion of PtdIns(3,5)P_2_ (ref^41^), resulted in a stronger block of both STING and TBK1 phosphorylation in response to either cGAMP delivery (**Fig. 2e**, right) or HT-DNA transfection (**Extended Data Fig. 4a**). Similar effects of YM201636 were observed at different time points after cGAMP delivery (**Extended Data Fig. 4b**) and in other cell lines (L929 and MEF; **Extended Data Fig. 4c, d**). However, prolonged treatment with YM201636 did not affect TBK1 activation in the RNA-sensing pathway (**Extended Data Fig. 4e**). These data, together with the PIKfyve depletion experiments, suggest that the kinase activity of PIKfyve, i.e., PtdIns(3,5)P_2_ production, is essential for STING-mediated TBK1 activation.

### I*n vitro* reconstitution of PtdIns(3,5)P_2_-mediated STING activation

Higher-order oligomerization of the STING dimers is a key mechanism for TBK1 activation^16,42^. We reasoned that the perinuclear clusters of STING vesicles might achieve signaling activation by binding to PtdIns(3,5)P_2_ and potentially other lipids, which promote STING oligomerization. Besides the PIKfyve product PtdIns(3,5)P_2_, activated STING might also access PtdIns(4)P and PtdIns(3)P, which are highly enriched in TGN and endosomes, respectively^26–28^. To identify which lipid(s) might be involved in STING activation, we developed an in vitro system to reconstitute STING-mediated TBK1 activation using cell-derived liposomes. In this system, various soluble, negatively charged short chain (diC8) phospholipids were directly added to the reactions (**Fig. 3a**), which are known to insert into membranes with well-characterized partition behaviors^43^. With 25 μM of soluble diC8 phosphoinositides, the membranes in our system were estimated to reach a final concentration of 0.5-1 mol% of phosphoinositides (see methods). When tested individually in this assay, PS and most phosphoinositide species showed limited effects, but PtdIns(3,5)P_2_ strongly stimulated cGAMP-induced phosphorylation of both STING and TBK1 (**Fig. 3b, c**, **Extended Data Fig. 5a**), leading to an electrophoretic mobility shift of STING indicative of its phosphorylation. Importantly, PtdIns(3,5)P_2_ sensitized STING activation in response to as low as 10 nM cGAMP (**Fig. 3d, e**). This in vitro, cell-free signaling activation required the presence of ATP, cGAMP, and PtdIns(3,5)P_2_ (**Extended Data Fig. 5b**) as well as STING (**Extended Data Fig. 5c**). The reaction produced similar results using liposomes derived from BJ cells overexpressing STING (**Extended Data Fig. 5d**). Together, our *in vitro* assay identified PtdIns(3,5)P_2_ as an enhancer of cGAMP-stimulated, STING-dependent TBK1 activation, consistent with the requirement of PIKfyve and its kinase activity in STING activation (**Fig. 2**).

**Figure 3.**
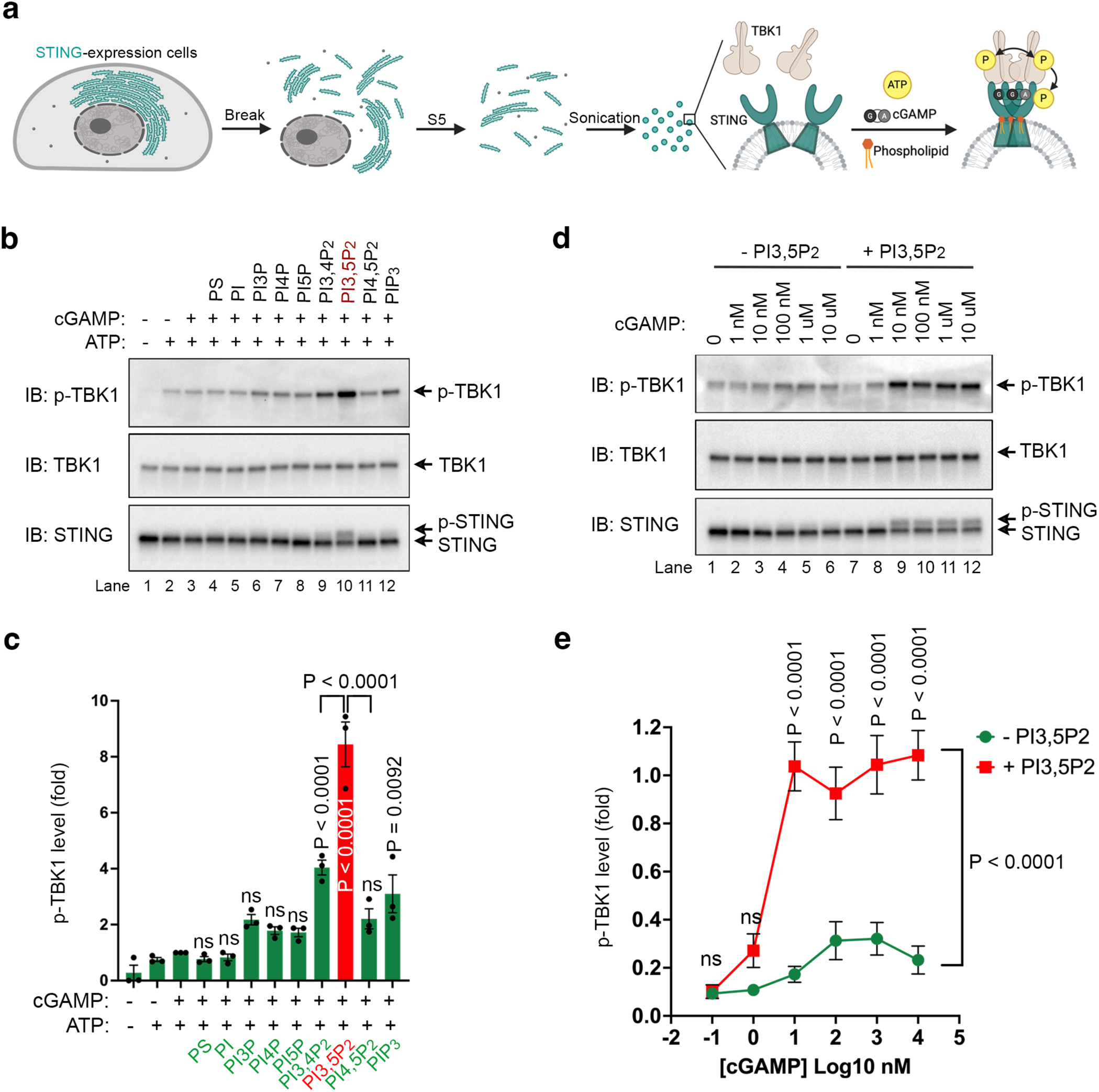
In *vitro* reconstitution identifies PtdIns(3,5)P_2_ as an enhancer of STING-mediated TBK1 activation. **(a)** Schematic diagram of STING signaling reconstitution using cell-derived liposomes. **(b)** The effects of individual phospholipid (25 μM in solution) on cGAMP-stimulated phosphorylation of TBK1 and STING on cell-derived liposomes in vitro. **(c)** Quantification of p-TBK1 levels in (b); mean ± SEM; n = 3. Two-way ANNOVA, followed by Turkey’s multiple comparisons tests. **(d)** PtdIns(3,5)P_2_ (25 μM in solution) sensitizes reconstituted STING and TBK1 phosphorylation to 10 nM of cGAMP. **(e)** Quantification of p-TBK1 levels in (d); mean ± SEM; n ≥ 4. Two-way ANNOVA, followed by Turkey’s multiple comparisons tests.

### PtdIns(3,5)P_2_ binds to STING

Phosphoinositide kinases often associate with phosphoinositide effectors^44^. Since PIKfyve associates with STING and PtdIns(3,5)P_2_ promotes STING signaling, we hypothesized that STING is a PtdIns(3,5)P_2_-binding protein. A purified C-terminal fragment (CT) of STING showed no phosphoinositide binding in PIP strips assays (**Extended Data Fig. 6a**), suggesting potential requirement of its transmembrane domain (TMD) in lipid binding. To test PtdIns(3,5)P_2_ binding to full length STING, a fluorescence resonance energy transfer (FRET) assay was developed, as previously reported in detecting the lipid binding to a transmembrane K^+^ channel^45^. Full-length human STING with a monomeric EGFP (mEGFP) fused to the C-terminus was used as the FRET donor and a soluble fluorescently labeled PtdIns(3,5)P_2_ [BODIPY TMR-PtdIns(3,5)P_2_] as the acceptor (**Fig. 4a**). The optimal excitation wavelength for the FRET experiment was determined to be ∼460 nm, which effectively excited EGFP with minimal cross-excitation of BODIPY TMR (**Fig. 4b**). If BODIPY TMR-PtdIns(3,5)P_2_ binds to STING-mEGFP, FRET should occur, causing a decrease of donor emission at 520 nm (Emission 1, Em1) and an increase of acceptor emission at 574 nm (Em2, **Fig. 4b**).

**Figure 4.**
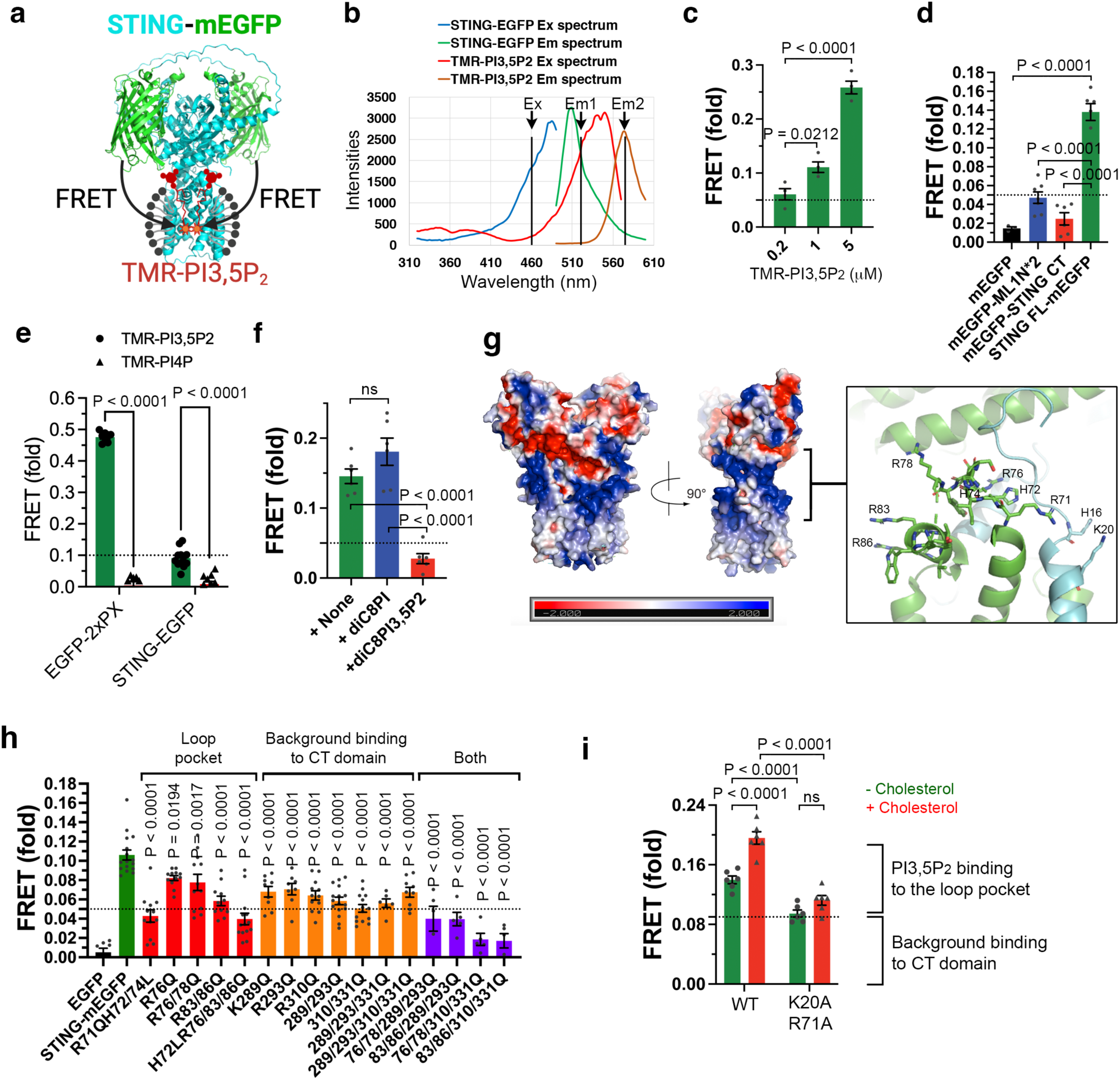
STING binds to PtdIns(3,5)P_2_ in vitro. **(a)** Schematic illustration of the FRET assay using STING-mEGFP and BODIPY® TMR- PtdIns(3,5)P_2_. **(b)** Excitation (Ex) and emission (Em) spectrum of STING-mEGFP and TMR-PtdIns(3,5)P_2_. Arrows point to the excitation (460 nm) and emission wavelengths (520 nm for EGFP and 574 nm for TMR) for the FRET assay. **(c)** FRET between various concentrations of TMR-PtdIns(3,5)P_2_ and 50 nM of STING-mEGFP. Unless otherwise indicated, FRET refers to the fraction of EGFP emission loss in the presence of TMR-PtdIns(3,5)P_2_. **(d)** FRET between 1 μM TMR-PtdIns(3,5)P_2_ and 50 nM mEGFP or indicated mEGFP fusion proteins. **(e)** FRET between 1 μM TMR-PtdIns(3,5)P_2_ or TMR-PtdIns(4)P and 50 nM EGFP-2xPX or STING-EGFP fusion proteins. **(f)** Competition FRET assay using unlabeled PtdIns(3,5)P_2_ or Ptdlns. 50 nM STING-mEGFP was mixed with 100 μM unlabeled PtdIns(3,5)P_2_ or Ptdlns before the addition of 1 μM TMR-PtdIns(3,5)P_2_. **(g)** Left: surface charge distribution on human STING dimer (PDB: 6NT5). Right: Expanded view of positively charged residues along the cytosolic loop of human STING. **(h)** FRET between 1 μM TMR-PtdIns(3,5)P_2_ and 50 nM STING-mEGFP or indicated mutants. **(i)** Comparing wild-type and the K20A/R71A PtdIns(3,5)P_2_-binding mutant of STING-mEGFP in FRET assays with 1 μM TMR-PtdIns(3,5)P_2_ ± 100 μM cholesterol. Data represent mean ± SEM; n = 4 for panel c; n ≥ 5 for all other panels. Two-way ANNOVA, followed by Turkey’s multiple comparisons tests.

With ∼50 nM of full length STING-mEGFP, a substantial decrease of EGFP emission was detected in the presence of BODIPY TMR-PtdIns(3,5)P_2_ in a dose-dependent manner (**Fig. 4c**). The observed FRET required full length STING, as mEGFP alone or its fusion protein with STING-CT 281-379 showed minimal emission reduction in the presence of BODIPY TMR-PtdIns(3,5)P_2_ (**Fig. 4d**). As a positive control for this assay, EGFP-2xPX, the recently reported high-affinity PtdIns(3,5)P_2_ probe derived from the amoeba *Dictyostelium discoideum*, showed dramatic FRET, with about 50% of EGFP quench upon the addition of BODIPY TMR-PtdIns(3,5)P_2_ (**Fig. 4e, Extended Data Fig. 6b**). A tandem repeat of the cytosolic N-terminal segment (ML1N*2) of transient receptor potential Mucolipin 1 (TRPML1) channel, which was previously shown to bind PtdIns(3,5)P_2_ (ref^41^), showed much weaker FRET with BODIPY TMR-PtdIns(3,5)P_2_ (**Fig. 4d**). In competition assays, unlabeled PtdIns(3,5)P_2_, but not PtdIns, abolished the FRET signal (**Fig. 4f**), indicating that the observed FRET was due to an interaction between PtdIns(3,5)P_2_ and STING. STING may encounter four phosphoinositide species during its trafficking form the ER through the Golgi complex to perinuclear endosomes: PtdIns(3)P, PtdIns(4)P, PtdIns(5)P, and PtdIns(3,5)P_2_ (ref^46,47^). Among them, PtdIns(3,5)P_2_ bound to STING with the highest affinity in competitive FRET assays (**Extended Data Fig. 6c**). In line with previous reports of the PI4KB/PtdIns(4)P signaling as a potential regulator of STING^48,49^, inhibitors of either PI4KB or PIKfyve similarly suppressed STING-TBK1 signaling (**Extended Data Fig. 6d, e**). However, No FRET signal was detected for either EGFP-2xPX or STING-EGFP when BODIPY TMR-PtdIns(3,5)P_2_ was replaced by BODIPY TMR-PtdIns(4)P (**Fig. 4e**), indicating that PtdIns(4)P does not directly bind to STING. Because acute depletion of the Golgi PtdIns(4)P blocks vesicle trafficking from the Golgi to peripheral compartments^50^, PtdIns(4)P may indirectly affects STING signaling through vesicle trafficking. This is consistent with a specific effect of PtdIns(3,5)P_2_, but none of the other phosphoinositide species, on STING-mediated TBK1 activation in vitro (**Fig. 3b, c**).

Since most of the detected PtdIns(3,5)P_2_ binding was reduced by unlabeled PtdIns(3,5)P_2_ but not PtdIns (**Fig. 4f**), the detected PtdIns(3,5)P_2_ binding in FRET may rely on electrostatic interactions between STING and the negatively charged headgroup of PtdIns(3,5)P_2_. To identify potential PtdIns(3,5)P_2_ binding sites on STING, we analyzed the surface charge distribution on the cryo-electron microscopy (EM) structure of full-length human STING dimer (PDB: 6nt5)^42^. This revealed positively charged regions right above the TMD, with most charges contributed by a cytosolic loop between the second and third transmembrane helices as well as by the N-terminal cytosolic tail from the other protomer (**Fig. 4g, Extended Data Fig. 7a**). Interestingly, various point mutations in the positively charged loop involving residues Arg71, His72, His74, Arg76, Arg78, Arg83, and Arg86 partially reduced STING binding to PtdIns(3,5)P_2_ (**Fig. 4h**), suggesting that this region may contribute to electrostatic interactions with PtdIns(3,5)P_2_. The partial reduction in PtdIns(3,5)P_2_-binding also suggests additional binding sites, possibly in the CT domain of STING. Mutations of various positively charged residues in the CT domain that is distant to the TMD domain caused a consistent ∼35% reduction in PtdIns(3,5)P_2_ binding in FRET assays (**Fig. 4h**). This suggests that the CT domain, which is away from the membrane in cells, may mediate basal PtdIns(3,5)P_2_-binding in detergent. Combination of point mutations in both the loop and the CT further reduced PtdIns(3,5)P_2_ binding (**Fig. 4h**). Thus, in the FRET-based lipid binding assays, STING bound to PtdIns(3,5)P_2_ through both the cytoplasmic loop region and the CT domain, the latter of which appeared to mediate background binding.

FRET is based on a transfer of energy between the donor and the acceptor, leading to reduced emission from the donor and increased emission from the acceptor. However, in FRET assays with 50 nM of STING-mEGFP and 1 μM of BODIPY TMR-PtdIns(3,5)P_2_, no obvious increase in BODIPY TMR emission was observed despite a consistent ∼10% loss of EGFP emission for wild type STING. This is likely due to the self-quenching of BODIPY TMR molecules when they were brought too close to a binding protein, as previously reported^51,52^. Consistently, purified full-length human STING without an mEGFP tag caused 3-4% emission loss of BODIPY TMR-PtdIns(3,5)P_2_ (**Extended Data Fig. 7b**), suggesting that STING might induce BODIPY TMR self-quenching by bringing multiple molecules of BODIPY TMR-PtdIns(3,5)P_2_ to close proximity. Thus, the energy transferred from STING-mEGFP (50 nM) might not be sufficient to overcome the self-quenching of BODIPY TMR (1 μM). After increasing STING-mEGFP to 500 nM and reducing BODIPY TMR-PtdIns(3,5)P_2_ to 250 nM, a consistent ∼5% increase of BODIPY TMR emission was observed, and such an increase was not observed in PtdIns(3,5)P_2_-binding mutants (**Extended Data Fig. 7c**). Thus, STING seems to bring PtdIns(3,5)P_2_ molecules into close proximity.

To understand how PtdIns(3,5)P_2_ binds to STING, we solved the cryo-EM structure of full length human STING in complex with PtdIns(3,5)P_2_ (Li et al., co-submitted). This revealed PtdIns(3,5)P_2_, together with a cholesterol analog cholesteryl hemisuccinate (CHS), as a molecular glue tethering STING dimers, thereby promoting cGAMP-induced STING oligomerization. In this structure, the phosphate group at the 5-position of the PtdIns(3,5)P_2_ inositol ring bound to a groove formed by three residues: H16 and K20 from the N-terminal tail of one STING protomer and R71 from the loop between the second and third transmembrane helices from another protomer (Li et al., co-submitted). Thus, mutations around the loop might have disrupted this pocket and thus reduced PtdIns(3,5)P_2_ binding in FRET assays (**Fig. 4g, h**). To test if the observed groove in cryo-EM is essential for PtdIns(3,5)P_2_ binding, we generated a K20A/R71A mutant. This mutant exhibited a reduction in PtdIns(3,5)P_2_ binding that was comparable to other loop mutants(**Fig. 4h, i**). Notably, wild-type STING, but not the K20A/R71A mutant, showed increased binding to PtdIns(3,5)P_2_ in the presence of cholesterol (**Fig. 4i**), consistent with a potential role of cholesterol in PtdIns(3,5)P_2_ binding to STING. These results indicate that STING directly binds to PtdIns(3,5)P_2_.

### PtdIns(3,5)P_2_-binding promotes STING trafficking from the ER

To investigate how PtdIns(3,5)P_2_ binding regulates STING signaling, we reconstituted wild-type (WT) STING or its PtdIns(3,5)P_2_-binding deficient mutant K20A/R71A in human U2OS cells which lack endogenous STING^18^. The WT STING showed extensive perinuclear trafficking upon cGAMP stimulation (**Fig. 5a**). However, the trafficking of the PtdIns(3,5)P_2_-binding deficient K20A/R71A mutant was severely impaired, with the vast majority of the mutant remaining on the ER, similar to an oligomerization-deficient STING mutant Q273A/A277Q (**Fig. 5a**). Consistently, while WT STING reconstituted TBK1 signaling in U2OS cells, the K20A/R71A mutant was largely defective (**Fig. 5b**), with the upregulation of type I interferon (*INFB1*) and interferon-stimulated genes (*CCL5* and *ISG20*) almost abolished (**Fig. 5c**). In vitro assays using liposomes derived from STING K20A/R71A-expressing cells showed no response to PtdIns(3,5)P_2_ in cGAMP-stimulated STING-TBK1 activation (**Fig. 5d, Extended Data Fig. 7d**). Besides interferon signaling, STING has primordial functions in noncanonical autophagy (macroautophagy-independent LC3 lipidation) and TFEB-mediated transcriptional upregulation of lysosomal biogenesis^20,53–58^, both of which were also reduced in cells expressing the K20A/R71A mutant of STING (**Fig. 5e, Extended Data Fig. 7e, f**). Thus, PtdIns(3,5)P_2_-binding supports both the canonical and noncanonical functions of STING.

**Figure 5.**
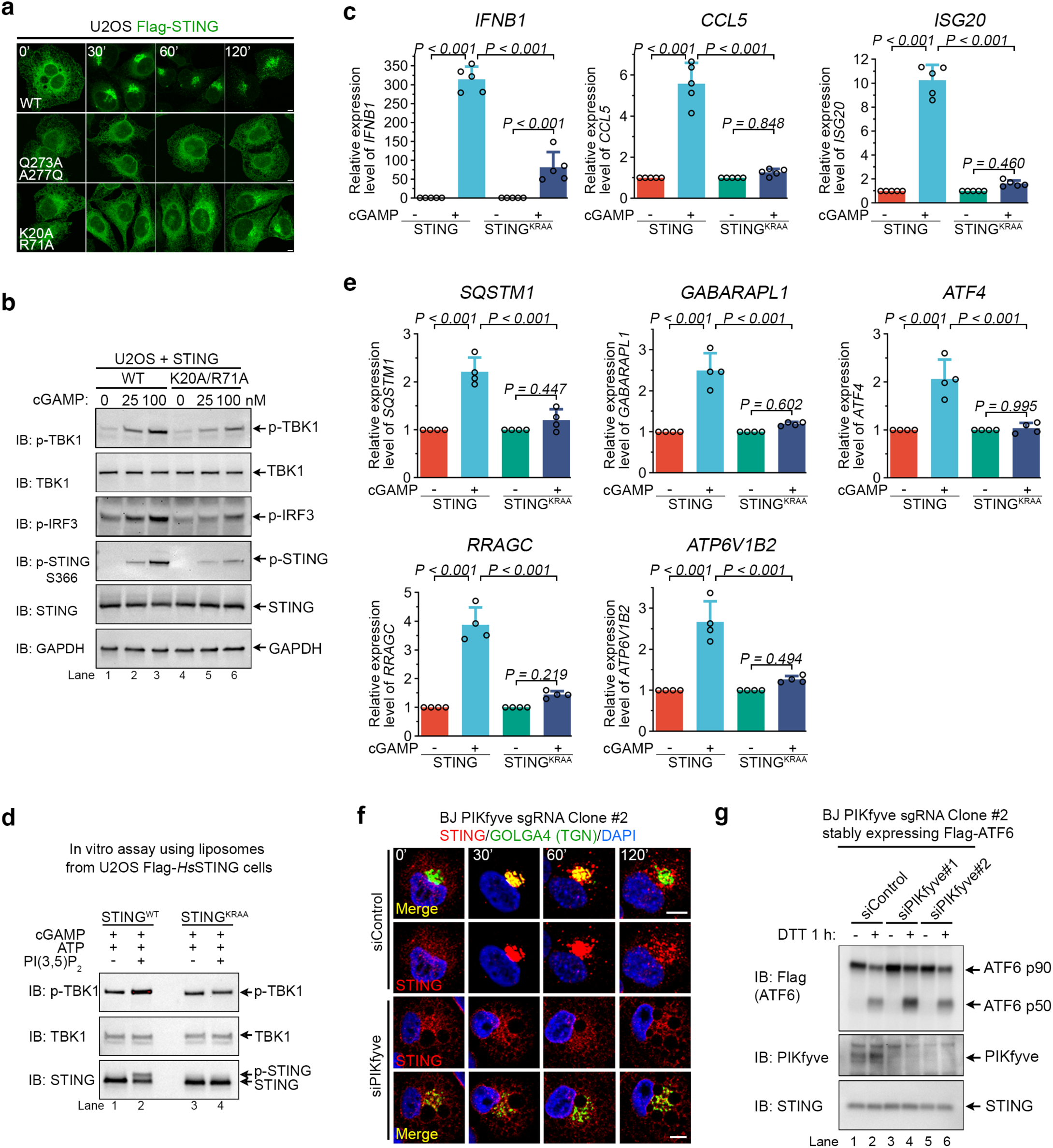
PtdIns(3,5)P_2_regulation of STING trafficking and signaling activation. **(a)** The PtdIns(3,5)P_2_-binding mutant (K20A/R71A) and an oligomerization-defective mutant (Q273A/A277Q) of STING have defects in trafficking from the ER to perinuclear clusters in BJ STING-KO cells. Cells were treated with or without 100 nM cGAMP and fixed at indicated time points for immuno-staining of Flag-STING. Bar, 5 μm. **(b)** The PtdIns(3,5)P_2_-binding mutant STING (K20A/R71A) shows reduced signaling capacity when reconstituted in U2OS cells which do not express endogenous STING. Cells were treated with 100 nM cGAMP and whole cell lysates were harvested 1 hour later for immunoblotting. **(c)** The PtdIns(3,5)P_2_-binding mutant of STING (K20A/R71A) has defects in upregulating interferon beta (*IFNB1*) and interferon stimulated genes (*CCL5* and *ISG20*). U2OS cells stably expressing STING or its mutant were treated with 200 nM cGAMP for 2 h, followed by RNA extraction and qRT-PCR. Mean ± sd; n = 5. Two-way ANNOVA, followed by Turkey’s multiple comparisons tests. **(d)** The PtdIns(3,5)P_2_-binding mutant of STING (K20A/R71A) does not respond to PtdIns(3,5)P_2_ (25 μM in solution) in in vitro reconstitution of STING-TBK1 signaling using U2OS-derived liposomes. **(e)** The PtdIns(3,5)P_2_-binding mutant of STING (K20A/R71A) has defects in upregulating TFEB target genes. U2OS cells stably expressing STING or STING mutant (K20A/R71A) were treated with 200 nM cGAMP for 6 h, followed by RNA extraction and qRT-PCR. Mean ± sd; n = 4. Two-way ANNOVA, followed by Turkey’s multiple comparisons tests. **(f)** Complete depletion of PIKfyve blocks STING trafficking in BJ cells. BJ PIKfyve CRISPR cells were transfected with nonspecific control or PIKfyve siRNA; three days later cells were treated with 100 nM cGAMP and fixed at indicated time points for co-staining of STING and GOLGA4, a TGN marker. **(g)** Complete depletion of PIKfyve does not affect ATF6 cleavage in BJ cells. BJ PIKfyve CRISPR cells stably expressing ATF6 were transfected control or PIKfyve siRNA; three days later cells were treated with or without 2 mM DTT for 1 hour and harvested for immunoblotting.

Given that STING oligomerization is essential for its trafficking from the ER^42^ and that PtdIns(3,5)P_2_-binding promotes STING oligomerization, our observation of a constitutive association between STING and PIKfyve likely indicates a role of PIKfyve on the ER in cGAMP-induced STING trafficking. Indeed, complete depletion of PIKfyve caused a severe block of STING trafficking from the ER (**Fig. 5f**). ATF6 is another ER-anchored transmembrane protein that upon ER stress traffics to the Golgi where it is proteolytically processed^59,60^. Complete depletion of PIKfyve did not affect ATF6 cleavage in response to dithiothreitol (DTT)-induced ER stress (**Fig. 5g**), indicating that trafficking of ATF6 from the ER to the Golgi does not require PtdIns(3,5)P_2_. The cargo selectivity for PIKfyve-regulated, ER-to-Golgi trafficking suggests that PIKfyve and its phospholipid product, PtdIns(3,5)P_2_, act on the cargo, but not the trafficking machineries, which is consistent with PtdIns(3,5)P_2_-promoted STING oligomerization.

### PtdIns(3,5)P_2_ enhances STING-dependent TBK1 autophosphorylation

Our in vitro signaling reconstitution indicates that the co-stimulation of cGAMP and PtdIns(3,5)P_2_ was sufficient for STING-dependent TBK1 activation (**Fig. 3**). However, the constitutive STING-PIKfyve association suggests that STING already has access to PtdIns(3,5)P_2_ on the ER. We thus further explored why STING cannot activate TBK1 when the Golgi is disassembled by Brefeldin A (**Fig. 1e**). We hypothesized that a relatively low level of PtdIns(3,5)P_2_ on the ER is sufficient for cGAMP-induced STING trafficking whereas higher PtdIns(3,5)P_2_ levels on post-Golgi endosomes are necessary for TBK1 activation by STING. This hypothesis is consistent with the enriched localization of PtdIns(3,5)P_2_ in endolysosomes instead of other organelles such as the ER^32,61,62^.

To test our hypothesis, we aimed to decouple STING trafficking from TBK1 activation through manipulating cellular PtdIns(3,5)P_2_ levels. Treating BJ cells with the PIKfyve inhibitor YM201636 for 18 h caused less severe endolysosomal vacuolation than complete PIKfyve depletion (**Extended Data Fig. 3b and 8a**), suggesting reduced but not complete removal of PtdIns(3,5)P_2_ in YM201636-treated cells. We further noticed that, while complete PIKfyve depletion severely blocked STING trafficking from the ER (**Fig. 5f**), pretreatment of BJ cells with YM201636 for 18 h did not affect cGAMP-stimulated STING trafficking to the perinuclear punctate compartments (**Extended Data Fig. 8b**). TBK1 recruitment to STING vesicles also appeared to be normal in YM201636-treated cells (**Extended Data Fig. 8b**), despite a clear defect in STING-dependent TBK1 activation in these cells (**Fig. 2e, Extended Data Fig. 4a-d**). Thus, the remaining levels of PtdIns(3,5)P_2_ in YM201636-treated cells was sufficient to support TBK1 recruitment and STING trafficking, but was insufficient for TBK1 activation.

To further validate our results, we aimed to decouple STING trafficking from TBK1 activation in the same pool of cells. One approach is to examine the trafficking of fluorescently labeled STING in live cells followed by cell lysis to probe TBK1 phosphorylation. Although the C-terminal fusion protein STING-mEGFP showed normal trafficking in response to cGAMP stimulation, it was defective in activating TBK1 (**Extended Data Fig. 8c**), likely due to a loss of TBK1 recruitment to the C-terminus of STING^16^. We thus took advantage of a recently developed signaling-competent STING-110mEGFP fusion protein in which mEGFP was inserted to the luminal loop of STING between the third and the fourth transmembrane helices^63^ (**Extended Data Fig. 8c**). BJ STING- KO cells stably expressing STING-110mEGFP showed normal activation of TBK1 signaling in response to cGAMP (**Extended Data Fig. 8c**). Pretreatment with the PIKfyve inhibitor YM201636 for 18 h did not affect the trafficking of STING-110mEGFP to perinuclear vesicle clusters but suppressed TBK1 activation in the same cells (**Extended Data Fig. 8d**), which is further confirmed by reduced p-TBK1 signals in immunofluorescence (**Extended Data Fig. 8e, f**). These results confirm that STING trafficking and TBK1 activation can be decoupled and that a higher threshold of PtdIns(3,5)P_2_ levels is likely required for STING-mediated TBK1 activation.

STING-mediated TBK1 activation requires high-order STING oligomerization which brings multiple TBK1 molecules into proximity allowing their trans-autophosphorylation at Ser172^16,42^. Consistently, the kinase dead mutant of TBK1, K38A, cannot be phosphorylated at Ser172 upon STING activation by cGAMP (**Extended Data Fig. 9a**), ruling out TBK1 phosphorylation and thus activation by other kinases. Although both STING trafficking and TBK1 recruitment appeared normal in YM201636-treated cells (**Extended Data Fig. 8b**), we hypothesized that such STING-TBK1 interaction was likely insufficient to trigger the trans-autophosphorylation of TBK1 due to low levels of PtdIns(3,5)P_2_ and reduced STING oligomerization. Indeed, when the STING-TBK1 interaction was evaluated by co-IP, a weaker interaction was detected in YM201636-treated cells (**Extended Data Fig. 9b**). Thus, TBK1 recruitment and activation can also be decoupled.

We further investigated the impact of PtdIns(3,5)P_2_ on TBK1 recruitment and activation. Following a typical in vitro reaction as shown in **Fig. 3a**, TBK1 was increased in the membrane fraction (**Extended Data Fig. 9c**), based on which we further developed an in vitro TBK1 membrane recruitment assay (**Fig. 6a**). TBK1 recruitment from the cytosol to STING-containing membranes was stimulated by cGAMP, which was not affected by the further addition of ATP or PtdIns(3,5)P_2_ (**Fig. 6b**). Thus, PtdIns(3,5)P_2_ activated the cGAMP/STING/TBK1 complex without affecting TBK1 recruitment. Consistent with normal TBK1 recruitment in PIKfyve-inhibited cells (**Extended Data Fig. 8b**), cGAMP-induced TBK1 recruitment to STING-containing membranes was also normal in BJ cells completely depleted of PIKfyve (**Fig. 6c**). However, when measured by co-IP, the STING-TBK1 interaction was almost abolished in the absence of PIKfyve (**Fig. 6d**) or when PIKfyve was inhibited (**Extended Data Fig. 9b**). Therefore, PtdIns(3,5)P_2_ is not essential for cGAMP binding or TBK1 recruitment to STING, but high levels of PtdIns(3,5)P_2_ is essential for strong interactions between STING and TBK1 for signaling activation (**Extended Data Fig. 10**). This is consistent with a model where PtdIns(3,5)P_2_-dependent high-order STING oligomerization facilitates TBK1 trans-autophosphorylation (**Fig. 6e**).

**Figure 6.**
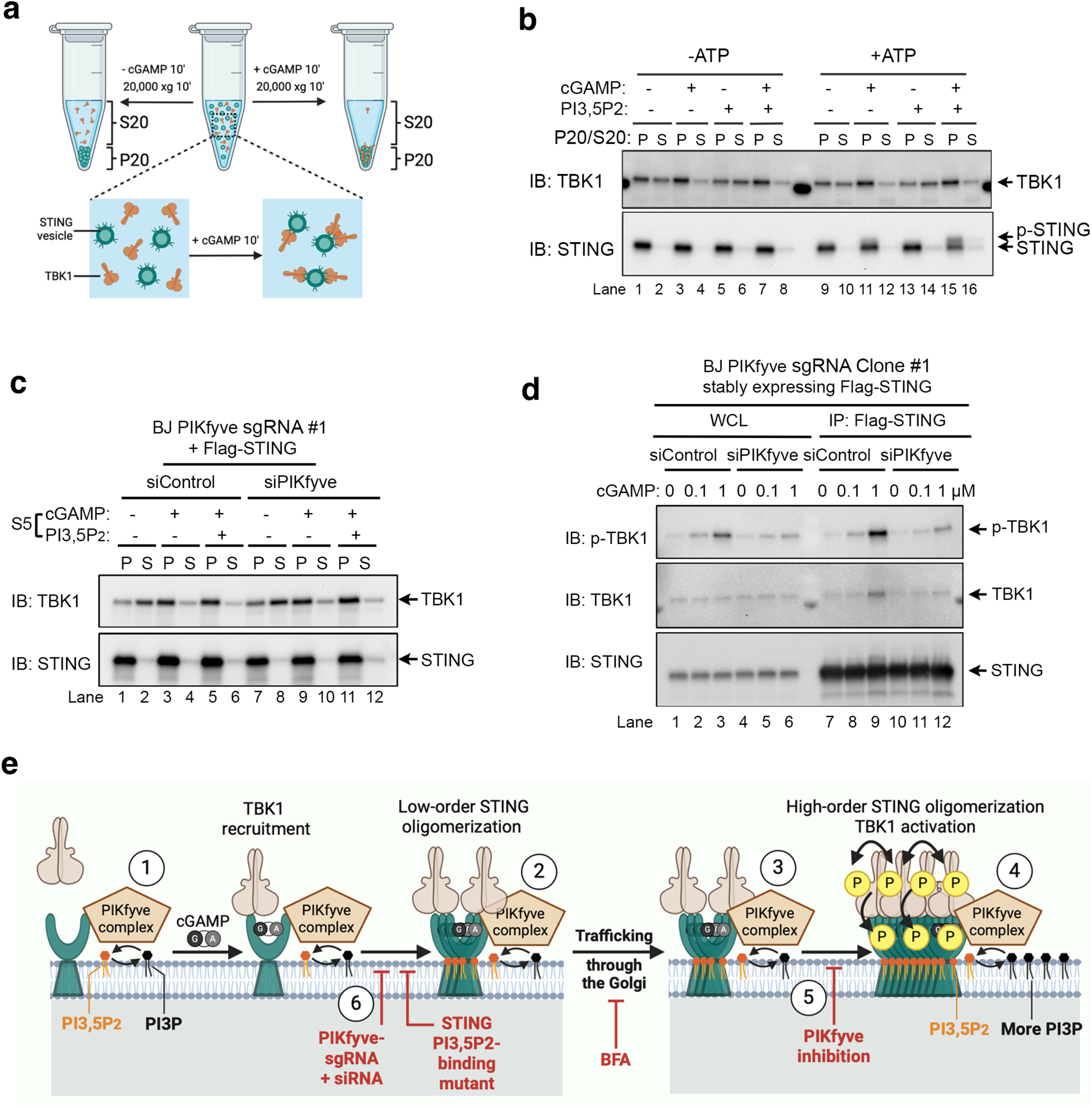
PtdIns(3,5)P_2_ romotes STING oligomerization to activate TBK1. **(a)** Schematic diagram of TBK1 membrane recruitment assay. **(b)** PtdIns(3,5)P_2_ does not affect cGAMP-induced TBK1 recruitment to STING-containing membranes. STING-containing liposomes derived from Flag-STING expressing COS7 cells were stimulated with 1 μM cGAMP at room temperature for 10 min in the presence or absence of 25 μM diC8 PtdIns(3,5)P_2_ in solution and the samples were then centrifuged at 20,000 *xg* for 10 min; pellets (P) and supernatants (S) were subjected to SDS-PAGE and immunoblotting. **(c)** Depletion of PIKfyve does not affect cGAMP-stimulated TBK1 recruitment to STING-containing membranes. STING-containing liposomes derived from Flag-STING expressing BJ PIKfyve CRISPR cells pretreated with control or PIKfyve siRNA for three days were stimulated and processed as in (b). The same concentrations of cGAMP and diC8 PtdIns(3,5)P_2_ as in (b) were used. **(d)** Complete PIKfvye depletion inhibits STING/TBK1 interaction in co-IP. BJ PIKfyve CRISPR cells stably expressing Flag-STING were pretreated with control or PIKfyve siRNA for three days and stimulated with cGAMP for 1 hour, followed by whole cell lysate harvest for IP with anti-Flag M2 affinity gel. **(e)** Schematic illustration of PtdIns(3,5)P_2_-regulated STING trafficking and TBK1 activation. (1) The PIKfyve complex constitutively associates with STING. (2) A low levels of PtdIns(3,5)P_2_ is maintained on the endoplasmic reticulum (ER) by PIKfyve which supports STING oligomerization and trafficking upon cGAMP stimulation. (3) Subcellular trafficking by itself does not activate STING signaling. (4) Higher PtdIns(3,5)P_2_ levels at TGN/post-Golgi endosomes, likely achieved by higher PI3P levels in these compartments, activates STING-mediated TBK1 autophosphorylation. (5) Partially reducing PtdIns(3,5)P_2_ levels by PIKfyve inhibition blocks TBK1 activation without affecting STING trafficking. (6) Completely removing PtdIns(3,5)P_2_ by PIKfyve depletion or mutating the PtdIns(3,5)P_2_-binding residues of STING abolishes STING trafficking from the ER and thus blocking downstream signaling activation. Note that, in in vitro assays using ER-derived STING-containing membranes, directly adding PtdIns(3,5)P_2_ can bypass the requirement of STING trafficking in TBK1 activation.

## DISCUSSION

Phosphoinositides are lipid messengers with strictly regulated subcellular distributions. The plasma membrane and TGN are enriched in PtdIns(4,5)P_2_ and PtdIns(4)P, respectively, and the signature phosphoinositides for endolysosomes are PtdIns(3)P and PtdIns(3,5)P_2_ (ref^46,47^). Studies on PtdIns(3,5)P_2_ have primarily focused on late endosomes and lysosomes, as disruption of PtdIns(3,5)P_2_ production causes severe endolysosomal vacuolation^40,62,64,65^. In addition to STING, PtdIns(3,5)P_2_ binds and activates multiple transmembrane proteins such as endolysosomal ion channels including transient receptor potential mucolipin channels (TRPML) ^66^ and mammalian two-pore channel proteins (TPCs)^67–69^. Besides endolysosomal regulation, a few studies also proposed roles for PtdIns(3,5)P_2_ in membrane trafficking between TGN and endosomes^61,70^ and in direct activation of ion channels on sarcoplasmic reticulum, a specialized type of ER in muscle cells^71,72^.

In this study, we identified PtdIns(3,5)P_2_ as an endogenous STING ligand which promotes both STING trafficking and signaling activation. The precise control of PtdIns(3,5)P_2_ regulation of STING is likely determined by the level of PtdIns(3,5)P_2_ on the local membrane. We propose that constitutive PIKfyve association with STING maintains a low level of PtdIns(3,5)P_2_ on the ER that binds to STING and ensures efficient STING oligomerization and trafficking upon cGAMP binding. This is in line with previously reported roles for PtdIns(3,5)P_2_ in the activation of ER-localized ion channels^71,72^. The low level of PtdIns(3,5)P_2_ on the ER may also support constitutive STING trafficking from the ER for lysosomal degradation, as PIKfyve-low expressing sgRNA clones showed much higher basal level of STING. A similar low threshold of PtdIns(3,5)P_2_ level in support of EGFR lysosomal degradation has been previously observed^65^. However, such low level of PtdIns(3,5)P_2_ on the ER is insufficient for TBK1 activation as blocking STING trafficking to the Golgi by Brefeldin A fully abolished TBK1 phosphorylation, despite normal TBK1 recruitment. PtdIns(3,5)P_2_ addition in our in vitro assays overcame the inability of cGAMP to activate TBK1 on ER-derived membranes, consistent with a requirement for higher levels of PtdIns(3,5)P_2_ in STING oligomerization and TBK1 activation. Accordingly, the subcellular localization of STING-mediated TBK1 activation was observed at TGN and post-Golgi endosomes, where PtdIns(3,5)P_2_ signaling was previously established^40,61,62,64,65,70^. The higher levels of PtdIns(3,5)P_2_ in perinuclear STING vesicles likely come from increased PI(3)P, the substrate used by PIKfyve to generate PtdIns(3,5)P_2_, on local membranes. The perinuclear STING vesicles are known to be positive for the early endosome marker RAB5 which is directly recruited by PI(3)P^73^. In addition to PtdIns(3,5)P_2_, the higher cholesterol levels at TGN and post-Golgi compartments may also contribute to STING activation, given the evidence that PtdIns(3,5)P_2_ cooperates with cholesterol in promoting STING oligomerization (Li et al., co-submitted). Other lipid modifications such as palmitoylation of STING in the Golgi have also been shown to promote STING oligomerization and downstream activation of TBK1 and IRF3^74,75^. However, PtdIns(4)P appears to be an indirect regulator of STING as it does not seem to bind to STING.

This study highlights a unique requirement of PtdIns(3,5)P_2_ for TBK1 activation in the DNA-sensing pathway but not the RNA-sensing pathway, indicating mechanistic differences in TBK1 activation in the two pathways. The requirement of TBK1 kinase activity for its own phosphorylation downstream of STING implies an autophosphorylation mechanism, which requires STING oligomerization^16^. Indeed, previous structural studies have ruled out the possibility of *cis*-autophosphorylation within the TBK1 dimer and suggested that higher-order oligomerization of TBK1 stimulates its *trans*-autophosphorylation^76–79^. Thus, PtdIns(3,5)P_2_-promoted STING oligomerization appears to be a key mechanism allowing for TBK1 *trans*-autophosphorylation specifically in the DNA-sensing pathway.

In summary, we identified PtdIns(3,5)P_2_ as an essential component of the cGAS-STING signaling pathway. Through constitutive association with STING, the PtdIns(3,5)P_2_-producing PIKfyve complex controls both STING trafficking from the ER and its activation of TBK1 in post-Golgi vesicles. This additional layer of regulation on STING oligomerization, trafficking, and signaling activation suggests new approaches for developing agonists or antagonists of STING to treat various human diseases.

## ACKNOWLEDGMENTS

We thank all members of the Chen and Tan groups for comments and discussions. We thank Conggang Zhang for the STING-110mEGFP construct and Peiqing Shi for the TBK1/IKKε knockout cell line. This work was supported by grants from the National Institutes of Health (R01-AI093967 to Z.J.C., R01CA273595 to X.-C.B. and X.Z. R01CA299257 to X.,C.B, X.Z and Z.J.C), the Welch Foundation (I-1389 to Z.J.C, I-1944 to X.-C.B. and I-1702 to X.Z.), and the Cancer Grand Challenge (CGCFUL-2021\100007) with support from Cancer Research UK and the US National Cancer Institute (to Z.J.C). J.X.T. acknowledges Cancer Research Institute Irvington Postdoctoral Fellowship and the National Institute of General Medical Sciences of the National Institutes of Health (NIH) award R35GM150506. Z.J.C. is a Howard Hughes Medical Institute (HHMI) investigator.

## AUTHOR CONTRIBUTIONS

J.X.T. conceived, designed and performed most experiments and analyzed data before the manuscript submission; B.L. performed all experiments and analyzed data during the revision, except in vitro assays which were performed by J.X.T.; J.L., X.Z. and X.-C.B. contributed to the design of STING mutations; T.L. and X.C. performed mass spectrometry experiments and data analysis; J.X.T. and Z.J.C. supervised the work; J.X.T., Z.J.C., X.Z. and X.-C.B. obtained funding; J.X.T. wrote the manuscript; T.L., B.L. and Z.J.C. edited the manuscript; all authors read and approved the manuscript.

## CONFLICT OF INTEREST

The authors declare no conflict of interest in this study.

## Notes

### Competing Interest Statement

The authors have declared no competing interest.

