## Supplementary information for "PtdIns(3,5)P_2_ Is an Endogenous Ligand of STING in Innate Immune Signaling"

#### METHODS

##### Resources Table

| REAGENT or RESOURCE | SOURCE | IDENTIFIER |
|---|---|---|
| **Antibodies** | | |
| Mouse monoclonal ANTI-FLAG® M2 | Sigma-Aldrich | Cat#F1804; IF 1:1000 |
| Mouse monoclonal ANTI-FLAG® M2 Affinity Gel | Sigma-Aldrich | Cat#A2220 |
| Rabbit monoclonal anti-STING | Cell Signaling Technology | Cat#13647; WB 1: 1000 |
| Rabbit monoclonal anti-p-STING(Ser366) (E9A9K) | Cell Signaling Technology | Cat#50907S; WB 1:1000 |
| Rabbit monoclonal anti-phospho-TBK1 Ser172 | Cell Signaling Technology | Cat#5483; WB 1: 1000 |
| Rabbit monoclonal anti-phospho-IRF3 Ser396 | Cell Signaling Technology | Cat#4947; WB 1: 1000 |
| Mouse monoclonal anti-TBK1/IKKi | Santa Cruz Biotechnology | Clone 72B587, Cat#sc-52931; WB 1:3000; IF 1: 500 |
| Rabbit polyclonal anti-STING | Tanaka et al.[25] | N/A. IF 1:2000 |
| Anti-GST HRP Conjugate | GE Healthcare | Cat#RPN1236; WB 1:5000 |
| Mouse monoclonal anti-β-Tubulin | Sigma-Aldrich | Cat#T8328; WB 1:5000 |
| Goat anti-rabbit IgG, HRP-linked | Cell Signaling Technology | Cat#7074; WB 1:1000 |
| Horse anti-mouse IgG, HRP-linked | Cell Signaling Technology | Cat#7076; WB 1:1000 |
| Monoclonal mouse anti-GM130 | BD Biosciences | Cat#610822; IF 1:1000 |
| Mouse anti-GOLGA4 | BD Biosciences | Cat#611280; IF 1:1000 |
| Rabbit anti-TGN38 | Santa Cruz Biotechnology | Clone #M290; IF 1:1000 |
| anti-LC3 | Proteintech | Cat#14600-1-AP; WB 1:1000 |
| anti-GABARAP | Proteintech | Cat#18723-1-AP; WB 1:1000 |
| anti-OSBP | Proteintech | Cat#11096-1-AP; WB 1:1000 |
| anti-TFEB | Bethyl Laboratories | Cat#A303-673A; WB 1:1000 |
| anti-GAPDH | Proteintech | Cat#60004-1-Ig; WB 1:1000 |
| **Bacterial and Virus Strains** | | |
| Stellar™ Competent Cells | TAKARA BIO INC. | Cat#636766 |

| | | |
|---|---|---|
| Sendai Virus (Cantell Strain) | Charles River Laboratories | Cat#10100816 |
| **Chemicals, Peptides, and Recombinant Proteins** | | |
| Phosphatidylinositol diC8 (PI diC8) | Echelon Biosciences | Cat#P0008 |
| Phosphatidylinositol 3-phosphate diC8 (PI(3)P diC8) | Echelon Biosciences | Cat#P3008 |
| Phosphatidylinositol 4-phosphate diC8 (PI(4)P diC8) | Echelon Biosciences | Cat#P4008 |
| Phosphatidylinositol 5-phosphate diC8 (PI(5)P diC8) | Echelon Biosciences | Cat#P5008 |
| Phosphatidylinositol 3,4-bisphosphate diC8 (PI(3,4)$P_2$ diC8) (P3408) | Echelon Biosciences | Cat#P3408 |
| Phosphatidylinositol 3,5-bisphosphate diC8 (PI(3,5)P2 diC8) | Echelon Biosciences | Cat#P3508 |
| Phosphatidylinositol 4,5-bisphosphate diC8 (PI(4,5)P2 diC8) | Echelon Biosciences | Cat#P4508 |
| Phosphatidylinositol 3,4,5-trisphosphate diC8 (PI(3,4,5)P3 diC8) | Echelon Biosciences | Cat#P3908 |
| BODIPY® TMR Phosphatidylinositol 3,5-bisphosphate | Echelon Biosciences | Cat#C-35M6 |
| BODIPY TMR Phosphatidylinositol 4-phosphate | Echelon Biosciences | Cat#C-04M6 |
| Lipofectamine 2000 | Thermo Fisher Scientific | Cat#11668019 |
| Lipofectamine RNAiMAX | Thermo Fisher Scientific | Cat#13778150 |
| Brefeldin A | Sigma-Aldrich | Cat#B5936 |
| PI4K beta inhibitor 3 | MedChemExpress LLC | Cat#HY-15679; CAS NO: 1245319-54-3 |
| diABZI (Compound 3) | Selleckchem | Cat#S8796; CAS NO: 2138498-18-5 |
| digitonin | Sigma-Aldrich | Cat#D141; CAS NO: 11024-24-1 |
| PIK-93 | MedChemExpress LLC | Cat#HY-12046; CAS NO: 593960-11-3 |
| Torin 1 | Cayman Chemica | Cat#10997; CAS NO: 1222998-36-8 |
| Avasimibe | Sigma-Aldrich | Cat#PZ0190; CAS NO: 166518-60-1 |
| IPTG | Sigma-Aldrich | Cat#I6758; CAS NO: 367-93-1 |
| 2’,3’-cGAMP | CHEMIETEK | Cat#CT-CGMAP; CAS NO: 1441190-66-4 |
| Puromycin dihydrochloride | Sigma-Aldrich | Cat#P9620; CAS NO: 58-58-2 |
| Blasticidin | InvivoGen | Cat#ant-bl-05; CAS NO: 3513-03-9 |
| Herring testis DNA (HT-DNA) | Sigma-Aldrich | #D6898 |
| YM201636 | Cayman Chemical | Cat#13576; CAS NO: 438545-06-3 |

| | | |
|---|---|---|
| **Deposited Data** | | |
| Apo STING dimer | Shang et al.[42] | PDB: 6nt5 |
| **Experimental Models: Cell Lines** | | |
| BJ-5TA | ATCC | CRL-4001 |
| BJ-5TA Flag-STING cells | This paper | N/A |
| BJ-5TA Flag-PIKfyve cells | This paper | N/A |
| BJ-5TA STING$^{-/-}$ cells | Gui et al.[54] | N/A |
| BJ-5TA STING$^{-/-}$ cells reconstituted with Flag-STING WT or mutants | This paper | N/A |
| BJ-5TA STING$^{-/-}$ cells reconstituted with STING-mEGFP or STING-110mEGFP, in the latter of which mEGFP was inserted into the second luminal loop after amino acid 110 | This paper | N/A |
| BJ-5TA TBK1/IKKε double knockout cells | This paper | N/A |
| BJ-5TA TBK1/IKKε double knockout cells reconstituted with TBK1 WT or K38A | This paper | N/A |
| BJ-5TA PIKfyve CRISPR clone #1 | This paper | N/A |
| BJ-5TA PIKfyve CRISPR clone #2 | This paper | N/A |
| BJ-5TA PIKfyve CRISPR clone #1 stably expressing Flag-STING | This paper | N/A |
| HEK293T | ATCC | CRL-3216 |
| COS7 | ATCC | CRL-1651 |
| COS7 Flag-STING cells | This paper | N/A |
| COS7 Flag-STING (K20A/R71A) cells | This paper | N/A |
| U2OS | ATCC | HTB-96 |
| U2OS Flag-STING cells | This paper | N/A |
| U2OS Flag-STING (K20A/R71A) cells | This paper | N/A |
| L929 | ATCC | CCL-1 |
| **Oligonucleotides** | | |
| Primer, siRNA, and gRNA sequences | This paper | Table S1 |
| **Plasmids** | | |
| pTY-Flag-STING (human) | This paper | N/A |
| pTY-Flag-STING-R71Q/H72/74L | This paper | N/A |
| pTY-Flag-STING-R76/78Q | This paper | N/A |
| pTY-Flag-STING-R83/86Q | This paper | N/A |
| pTY-Flag-STING-R293Q | This paper | N/A |
| pTY-Flag-STING-R310Q | This paper | N/A |
| pTY-Flag-STING-R289/293Q | This paper | N/A |
| pTY-Flag-STING-R310/331Q | This paper | N/A |
| pTY-Flag-STING-R293/310/331Q | This paper | N/A |
| pCDH-Flag-STING-WT (human) | Xun, et al.[18] | N/A |

| | | |
|---|---|---|
| pCDH-Flag-STING-K20A/R71A | This paper | N/A |
| pCDH-Flag-STING-Q273A/A277Q | This paper | N/A |
| pCDH-PIKfyve | This paper | N/A |
| pCDH-Flag-PIKfyve | This paper | N/A |
| pCDH-V5-PIKfyve | This paper | N/A |
| pCDH-GFP-PIKfyve | This paper | N/A |
| pCDH-PIKfyve (K1832E) | This paper | N/A |
| pCDH-PIKfyve (sgRNA resistant, siRNA resistant) | This paper | N/A |
| pCDH-Flag-PIKfyve (sgRNA resistant, siRNA resistant) | This paper | N/A |
| pCDH-PIKfyve (K1832E, sgRNA resistant, siRNA resistant) | This paper | N/A |
| pCDH-Flag-PIKfyve (K1832E, sgRNA resistant, siRNA resistant) | This paper | N/A |
| pJV0047 | Vines et al.[80] | Addgene #205139 |
| pCDH-EGFP-2xPX | This paper | N/A |
| pTY-TBK1-Flag | This paper | N/A |
| pTY-TBK1-K38A-Flag | This paper | N/A |
| pET28b-mEGFP-ML1N*2 | This paper | N/A |
| pET28b-mEGFP-STING 281-379 | This paper | N/A |
| pCDNA3-mEGFP-Flag | This paper | N/A |
| pCDNA3-Flag-STING-mEGFP | This paper | N/A |
| pCDNA3-Flag-STING-mEGFP mutants (K289Q; R293Q; R310Q; 289/293Q; 310/331Q; 289/293/331Q; 289/293/310/331Q; R71Q/H72/74L; R76/78Q; R83/86Q; H72L/R76/83/86Q; 76/78/289/293Q; 83/86/289/293Q; 76/78/310/331Q; 83/86/310/331Q, K20A/R71A) | This paper | N/A |
| pCS-STING-mEGFP | Han, et al.[63] | N/A |
| pCS-STING-110mEGFP | Han, et al.[63] | N/A |
| **Software and algorithms** | | |
| Adobe Photoshop 25.9.1 | Adobe | https://www.adobe.com/ |
| BioRender | BioRender | https://www.biorender.com |
| PyMOL Molecular Graphics System | Schrödinger | https://www.pymol.org/ |
| Leica Application Suite X 3.5.5.19976 | Leica Microsystems | https://www.leica-microsystems.com/ |
| jamovi 2.6.44.0 | The jamovi project | https://www.jamovi.org/ |
| TBtools-II | Chengjie Chen | https://github.com/CJ-Chen/TBtools-II |
| ImageJ | National Institutes of Health | https://imagej.net/ij/ |
| Microsoft Office Excel 2016 | Microsoft | https://www.office.com/ |

| | | |
|---|---|---|
| Microsoft Office Power Point Professional Plus 2016 | Microsoft | https://www.office.com/ |
| GraphPad Software 9.0.0 | GraphPad Software | GraphPad Software |

#### METHOD DETAILS

##### Cell culture, transfection, and treatments

BJ-5TA, HEK293T, Cos7, L929, U2OS, and MEF cells were cultured in Dulbecco's Modified Eagle's Medium (DMEM, Corning) supplemented with 10% (v/v) FBS and 1× Antibiotic-Antimycotic (amphotericin B, penicillin, streptomycin) at 37 °C in an atmosphere of 5% (v/v) $CO_2$. Unless otherwise indicated, DNA was transfected using Lipofectamine 2000 (Thermo Fisher Scientific) following manufacturer's instructions. Transfection of siRNA was based on Lipofectamine RNAiMax (Thermo Fisher Scientific) following manufacturer's instructions. Cell delivery of cGAMP was achieved by digitonin-mediated permeabilization[17]. cGAMP was mixed in digitonin permeabilization buffer (50 mM HEPES pH 7.0, 100 mM KCl, 3 mM $MgCl_2$, 0.1 mM DTT, 85 mM Sucrose, 0.2% BSA, 1 mM ATP, 8 μg/mL Digitonin) which was warmed to 37 °C right before use. Cell culture media were replaced with digitonin buffer containing cGAMP and permeabilization was allowed to go for 10 min before changing back to normal media. For small molecule inhibitor treatment, cells were grown into 90% confluency and treated with indicated inhibitor for 1 hour before digitonin-mediated cGAMP delivery or treated at the same time of HT-DNA transfection or virus infection because DNA or virus takes hours to activate STING signaling. For purification of proteins from HEK293T cells, transfection was performed using polyethylenimine (PEI) with a DNA:PEI ration of 1:3 and cells were harvested for protein purification 24-48 hours post transfection.

##### Molecular cloning and cell line generation

STING C terminal coding sequences were cloned into pET42b for bacterial expression of GST-tagged fragments. Point mutations were generated by PCR using primers carrying target mutations, followed by DPN1-mediated digestion of DNA templates. Human TBK1 open reading frame was cloned into pCDNA3 for mutagenesis manipulations. To allow for TBK1 re-expression in TBK1/IKKε double knockout BJ cells, sgRNA resistant mutations were introduced to TBK1 cDNA by site-directed mutations through polymerase chain reaction (PCR). For stable expression of C terminally Flag-tagged TBK1 and its mutants or N terminally Flag-tagged human STING, the relative DNA sequences were subcloned into pTY lentiviral vector. For viral packaging in one well from a 6-well plate, the pTY vector, pMD2.G and psPAX2 (Addgene, #12259 and #12260, from Dr. Didier Trono, École Polytechnique

Fédérale de Lausanne, Lausanne, Switzerland) were transfected into HEK293T cells in a ratio of 4:3:1 (1 μg, 0.75 μg, 0.25 μg, respectively). To generate cell lines stably expressing a target protein, cells were infected with indicated lentiviruses in the presence of 10 μg/ml polybrene, followed by chemical selection (puromycin or blasticidin) for at least 7days. For all cell types, puromycin was used at a final concentration of 2 μg/ml; For BJ cells blasticidin was also used at 2 μg/ml. When generating the BJ TBK1/IKKε double knockout cell line, all the resistant markers were used. Thus, re-expression of TBK1 was achieved without chemical selection, but after re-expression cells were examined by immunofluorescence to make sure the majority of cells were expressing TBK1 or its mutant.

**CRISPR knockout cell lines**

The sgRNA targeting sequences (Table S1) were ligated into plentiCRISPR v2 (Addgene Plasmid #52961). Packaging of CRISPR viruses was performed 6-well plates similarly to pTY lentivial packaging described above. Cells were infected with CRISPR lentiviruses also in 6-well plates for 48 hours in the presence of 10 μg/ml polybrene before selected with puromycin (2 μg/ml). One week after selection, the CRISPR pool were analyzed by western blot to determine knockout efficiency of specific sgRNAs. The pools generated from sgRNAs with highest knockout efficiency were chosen for colonization of knockout cells. Monoclones were tested by western blotting, followed by DNA sequencing to confirm gene knockout.

**Immunoprecipitation and Immunoblotting**

Cells were lysed with immunoprecipitation (IP) buffer (50 mM HEPES, pH 7.5; 100 mM NaCl; 5 mM $MgCl_2$; 5 mM NaF; 2 mM $Na_3VO_4$; 0.5% Triton X-100; protease inhibitor cocktail tablet) and lysed completely on ice. Lysates were centrifuged at 20,000 ×g for 5 min and supernatants collected as whole cell lysates. For immunoblotting, whole cell lysates were mixed with equal volumes of 2× sodium dodecyl sulfate (SDS) loading buffer (100 mM Tris-HCl, pH 6.8; 4% SDS; 0.2% bromophenol blue; 20% glycerol; 2% 2-mecaptoethonol) and heated for 5-10 min at 95 °C before loaded for SDS-polyacrylamide gel electrophoresis (SDS-PAGE). For immunoprecipitation, whole cell lysates were incubated with M2 beads for 2 hours at 4°C, followed by washing with IP buffer three times. For each washing, beads were resuspended in 500 μl IP buffer and spin at 3,500 ×*g* for 10 seconds; all buffer was completely removed using a gel loading tip. Protein complexes bound on beads were eluted by adding 2 × SDS loading buffer and heated at 95 °C for 10 min. The co-IP'ed proteins were analyzed by SDS-PAGE and immunoblotting.

**Immunofluorescence Microscopy**

Human BJ Cells were seeded on covered slips in a 24-well plate. 24 hours later, cells were treated as indicated and washed with phosphate-buffered saline (PBS) twice before fixation with 4% polyformaldehyde (in PBS) for 5 min at room temperature. Cells were permeabilized with 0.25% Triton X-100 in PBS for 5 min at room temperature and blocked with 3% BSA in PBS for 1 hour at room temperature. Cells on coverslips were then incubated with primary antibodies at 4 °C overnight or at 37 °C for 2 hours. Cells were washed with PBS twice and incubated with Alexa-488 and/or -555 conjugated secondary antibodies for 1 hour at room temperature. Cell coverslips were mounted onto microscope slides using mounting media containing 4′,6-diamidino-2-phenylindole (DAPI) for nuclear staining (Vector Laboratories, #H-1200) and fixed with nail coat. Slides were examined and images collected with a Nikon A1R confocal microscope. All images in one experiment were taken with the same software setting and further assembled and equally processed in Adobe Photoshop.

**RNA isolation and real time qRT-PCR**

Total RNA was extracted from cells cultured in 6-well plates using the PureLink RNA Mini Kit (Thermo Scientific). Cells were washed twice with ice-cold PBS and lysed in 300 µl Lysis Buffer supplemented with 1% β-mercaptoethanol. Homogenization was performed by passing the lysate through a 21-gauge syringe needle five times, followed by addition of an equal volume of 70% ethanol. The mixture was vortexed and loaded onto a spin cartridge, then centrifuged at 12,000 ×g for 15 seconds at room temperature (RT). To eliminate genomic DNA contamination, an on-column DNase digestion step was implemented with PureLink DNase (Thermo Scientific): 80 µl DNase Solution (8 µl 10× Buffer, 10 µl DNase, 62 µl RNase-free water) was applied to the membrane and incubated for 15 minutes at RT. The cartridge was sequentially washed with 350 µl Wash Buffer I and two aliquots of 500 µl Wash Buffer II. After membrane drying by centrifugation (12,000 ×g, 1 minute), purified RNA was eluted in 30 µl RNase-free water and stored at −80°C. RNA integrity and concentration were verified spectrophotometrically with NanoDrop (Thermo Scientific). Reverse transcription was performed using the iScript cDNA Synthesis Kit (Bio-Rad) in a 20 µl reaction volume containing 1 µg RNA, 4 µl 5× iScript Reaction Mix, and 1 µl iScript Reverse Transcriptase. The thermal profile included: priming at 25°C for 5 minutes, cDNA synthesis at 46°C for 20 minutes, and enzyme inactivation at 95°C for 1 minute. Samples were held at 4°C for short-term storage. qRT-PCR assays were conducted with SYBR Green Universal Master Mix (Thermo Scientific) on a QuantStudio 3 thermocycler (Thermo Scientific). Each 20 μl reaction system contained 10 μl SYBR Green Mix, 1 μl cDNA template, 0.5 μl each of forward and reverse primers (10 μM), and 8 μl nuclease-free water. Primers were sourced from PrimerBank or GETPrime databases, or designed via Primer3Plus with the following criteria: amplicon length 80–200 bp, GC content 40–60%, and avoidance of secondary structures or repetitive sequences. The cycling protocol included an initial denaturation (95°C, 10 minutes), followed by 40 cycles of denaturation (95°C, 15 seconds) and annealing/extension (60°C, 1 minute), with a final melt curve analysis. All experiments incorporated three biological replicates (independent cell cultures) and three technical replicates (per cDNA sample). Gene expression was quantified using the $2^{-\Delta\Delta Ct}$ method via quickQrtPCR/TBtools-II software[81].

##### *In vitro* reconstitution of STING signaling activation in cell-derived liposomes

The cGAMP-stimulated STING signaling was found at perinuclear compartments where STING was heavily accumulated. Because directional vesicle trafficking and accumulation of STING was disrupted in cell-derived liposomes, a high STING concentration in this system was achieved by STING overexpression. Cells stably expressing high levels of Flag-tagged full length human STING were cultured in 15-cm dishes to 100% confluency. Cells were then used to generated liposomes containing STING in its native topology. Because both cholesterol and $PI(3,5)P_2$ bind to STING to facilitate oligomerization and signaling activation, the response to $PI(3,5)P_2$ in the in vitro assay required a certain level of cholesterol on ER-derived STING-containing liposomes. When U2OS cells were used for this assay, it required pretreatment with avasimibe to increase basal cholesterol levels on the ER.

Cells were washed with cold assay buffer (20 mM HEPES, pH 7.5, 250 mM sorbitol, 10 mM $MgCl_2$, 5 mM NaF, 2 mM $Na_3VO_4$) twice and all buffer was completely removed. An additional 500 μl cold assay buffer containing protease inhibitors was added to the dish and cells were scraped off and transferred into a 1.5 ml tube on ice. Samples were passed through a 27 G needle twice and centrifuged at 1000 × *g* for 3 min. Supernatants (S1) were moved into a new 1.5 ml tube and centrifuged at 5000 × *g* for 5 min. Supernatants (S5) were transferred into another new 1.5 ml tube. The S5 sample contained cell-derived membranes with full length STING and all components in the cytosol including TBK1. The S5 sample was diluted to a final protein concentration of 1-2 μg/μl, from which 20 μl was used for each *in vitro* reaction in a new 1.5 ml tube. The reaction tubes with diluted S5 samples were bath sonicated for 5 min at room temperature and cGAMP was added to a final concentration of 1 μM unless otherwise indicated. When addition of phospholipids was required, they can be added before or after sonication with similar effects. Upon cGAMP addition, the tubes were kept at room temperature for 8 min to allow for cGAMP binding and STING complex assembly. Then ATP was quickly added to a final concentration of 2 mM and the tubes were immediately transferred to a 37 °C heat block where the reaction was allowed to go for 10 min. Reaction was stopped by adding equal volumes of 2 × SDS loading buffer and samples were immediately heated at 95 °C for 5 min. The samples were analyzed by Western Blot.

Based on reported partitioning tests[43], diC8 $PI(4,5)P_2$ has an established partition behavior. If we assume diC8 $PI(3,5)P_2$ has a similar partitioning behavior to diC8 $PtdIns(4,5)P_2$, adding 25 uM of diC8 $PtdIns(3,5)P_2$ would result in a final mole fraction of 0.66% of $PtdIns(3,5)P_2$ in membranes. Below are the estimated calculations. We typically started with one 15-cm dish of COS7 cells which were estimated to have $1 \times 10^7$ cells with an average membrane area/cell = 20,000 μm². As each lipid occupies about 55 Å$^2$, total number of lipids per μm² (bilayer) would be about $3.6 \times 10^6$**.** Total lipid number from all the cells would be about $7.2 \times 10^{17}$, in a final volume of about 500 ul. Thus, the total lipid concentration would be around 2.4 mM. If the S5 fraction contained 20% of the lipids, then the total lipid concentration in S5 was about 480 uM. The most often dilution of S5 before final reaction is 1:2 which would give a final concentration of total membrane lipids of 240 uM.

Mass balance: $c_{total} = c_f + X_b \times c_L \times \gamma$

Total diC8 $PI(3,5)P_2$: $c_{total} = 25\,\mu M = 0.025\,mM$

Total accessible membrane lipids: $c_L \times \gamma = 120\,\mu M = 0.120\,mM$

$\gamma$ is the fraction of membrane lipids accessible to diC8 $PI(3,5)P_2$, which should be 0.5 as only the surface leaflet is accessible.

$X_b$ is the mol% of diC8 $PI(3,5)P_2$ after partitioning reaches an equilibration.

$c_f$ is the free diC8 $PI(3,5)P_2$ concentration after partitioning reaches an equilibration.

Polynomial for diC8-$PI(4,5)P_2$ from Collins et al.[43]:

$$log_{10}X_b = 0.00642252y^4+0.051329y^3+0.0644365y^2+0.685428y-1.07$$

where $y = log_{10}c_f$, and $c_f$ is in mM.

After equilibration,

Free diC8 $PI(3,5)P_2$ concentration ($cf$): 24.2 μM

Mol% of diC8 $PI(3,5)P_2$ in membrane (*Xb*): 0.0066 or 0.66%

**Estimation of the average molar fraction of PtdIns(3,5)$P_2$ in endolysosomal membranes**

The cellular $PI(3,5)P_2$ is about 0.04% out of all cellular phosphoinositol (PI) lipids[37]. As cells have about 10% PI lipids, $PI(3,5)P_2$ is about 0.004% of total cellular lipids. Most cellular PtdIns(3,5)$P_2$ is known to be enriched on endolysosomes (late endosomes and lysosomes). While the ER (~55%), mitochondria (~30%), and plasma membrane (~7.5%) together account for ~92.5% of total cellular lipids. The remaining 7.5% of lipids comes from all the remaining organelles including at least seven populations: lipid droplets, recycling endosomes, Golgi, peroxisomes, early endosomes, late endosomes, and lysosomes. Thus, late endosomes and lysosomes contribute only less than 2% of total cellular lipids, consistent with literature[82]. Thus, the average level of PtdIns(3,5)$P_2$ on total endolysosomes would be about 0.2%.

PtdIns(3,5)$P_2$ is enrichment in microdomains. Note that PtdIns(3,5)$P_2$ has strict turnover regulation, as the PIKfyve complex carries both the kinase and the FIG4 PtdIns(3,5)$P_2$-phosphatase. Thus, the local PtdIns(3,5)$P_2$ mol% would be significantly higher than the average 0.2% across endolysosomes. Indeed, PtdIns(3,5)$P_2$ is enriched in microdomains on endolysosomes[83]. Recent progress also revealed the enrichment of PtdIns(3,5)$P_2$ on a few selective endosomes[84], out of hundreds of endolysosomes per cells. Thus, the local PtdIns(3,5)$P_2$ concentrations on enriched membranes should be much higher than the estimated average of 0.2 mol%.

**TBK1 membrane recruitment assay**

TBK1 membrane recruitment assay was derived from the above in vitro signaling activation assay using cell derived STING-containing membranes. After 10 min of reaction at 37 °C, half of sample was saved as total sample and the other half was centrifuged at 20000 *xg* for 10 min at 4 °C. Pellet and supernatant were diluted with SDS loading buffer to the same volume of the total sample and analyzed by immunoblotting.

**Protein purification**

GST-tagged STING C terminus fragments were expressed in and purified from *E. coli*. Sequencing verified pET42b-STING-CT constructs were transformed into Rosetta competent cells (MilliporeSigma) and monoclones from kanamycin selection plates were expanded in 100 ml liquid LB media at 37 °C. When $OD_{600}$ of the culture reached

0.5, Isopropyl β-D-1-thiogalactopyranoside (IPTG) was added to a final concentration of 0.5 mM and the culture was allowed to incubate at 37 °C with shaking for 10 hours. Cells were pelleted by spin at 10, 000 × *g* for 10 min, resuspended in lysis buffer (50 mM HEPES, pH 7.5, 100 mM NaCl, 1% Triton X-100, protease inhibitor), sonicated and centrifuged at 20,000 × *g* for 10 min. Supernatants were collected and passed through a glutathione agarose (Thermo Fisher Scientific) column at 4 °C. Unbound proteins were washed off using washing buffer (20 mM Hepes, pH 7.5, 100 mM NaCl) and GST-STING-CT were eluted with elution buffer (20 mM Hepes, pH 7.5, 100 mM NaCl, 10 mM glutathione). Eluted protein was concentrated using Amicon Ultra centrifugal filter units with the pore size of 10 kDa (Sigma, #Z648027), which also removed glutathione. Protein concentration was determined by Bio-Rad Bradford Protein Assays and protein purity was assessed by gel staining.

Human STING open reading frame was fused with Flag-tag at its N-terminus and monomer EGFP (mEGFP A206K) at its C-terminus, and the fusion sequence was cloned into pCDNA3. Flag-STING-mEGFP was expressed in HEK293T cells by transient transfection with PEI. Each 15-cm dish with 75% confluency of HEK293T cells was transfected with 20 μg of DNA and cells were harvested 24 hours after transfection. Cells were washed and lysed in a buffer containing 20 mM Hepes, pH 7.5, 100 mM NaCl, 10 mM MgCl2, and 1% Triton X-100. Cell lysates were spin at 20,000X*g* for 5 min. Supernatant was used for immuno-purification of Flag-STING-mEGFP by anti-Flag M2 affinity gel, and the protein was eluted with Flag peptide in 20 mM Hepes, pH 7.5, 100 mM NaCl, and 0.1% Triton X-100. mEGFP-Flag was similarly purified from HEK293T cells. For purification of mEGFP-STING-C-tail, the relevant DNA sequences were cloned into pET28b for bacterial expression. All purified proteins were aliquoted immediately and stored at -80 °C.

The plasmid encoding the 6×His-EGFP-2×PX recombinant protein[80] (pJV0047, Addgene #205139) was verified by sequencing and transformed into E. coli BL21(DE3) competent cells (Thermo Fisher Scientific). A single colony selected on a kanamycin-containing plate was inoculated into 5 ml of LB broth and cultured overnight at 37°C with shaking. Subsequently, 1 ml of culture was transferred into 200 ml of fresh LB broth and grown at 37°C until the $OD_{600}$ reached 0.6–0.8. Protein expression was induced by adding IPTG to a final concentration of 0.2 mM, followed by incubation at 30°C for 6 hours. Cells were harvested by centrifugation at 5,000 × g for 15 min and resuspended in lysis buffer (50 mM HEPES pH 7.5, 300 mM NaCl, 1% Triton X-100, 1 mg/ml lysozyme, and protease inhibitor cocktail). After 30 min of incubation on ice, cells were sonicated and centrifuged at 12,000 × g for 15 min at 4°C. The supernatant was loaded onto a HisPur Ni-NTA Superflow Agarose column (Thermo Fisher Scientific #25214) pre-equilibrated with lysis buffer. Unbound proteins were removed by washing with 20 column volumes of wash buffer (50 mM HEPES pH 7.5, 300 mM NaCl, 20 mM imidazole). Bound His-tagged protein was eluted with elution buffer (50 mM HEPES pH 7.5, 300 mM NaCl, 500 mM imidazole). The eluate was concentrated using an Amicon Ultra-4 Centrifugal Filter Unit (30 kDa cutoff; Millipore Sigma #UFC803024). Protein concentration was determined by the Bradford assay (Bio-Rad) using bovine serum albumin (BSA) as a standard, and purity was assessed by SDS-PAGE followed by Coomassie blue staining.

**FRET-based lipid binding assay**

The FRET-based binding assay was carried out in a 50 μl reaction in 20 mM Hepes, pH 7.5, 100 mM NaCl, and 0.1% Triton X-100. Each reaction contains 100 nM EGFP fusion protein, 1 μM Bodipy-TMR-PtdIns(3,5)$P_2$, and various concentrations of competing phosphoinositides. When testing the impact of cholesterol on phosphoinositide binding, 100 μM $CaCl_2$ was included in the binding buffer. All components were first mixed well in a total of 50 μl/reaction in 1.5 ml tubes and then 40 μl from each tube was transferred (without causing bubbles) to a black 384-well plate. The samples were excited at 460 nm and emission was read at 520 nm and 574 nm, respectively.

**PIP strips**

GST-STING-CT 281-379 was purified from *E. coli*. for PIP strips assays. PIP strips membrane was blocked with 3% BSA in PBS-T (0.1% v/v Tween-20) for 1 hour at room temperature, followed by an additional 1-hour incubation with 0.2 μg/ml GST-PLCδ-PH or GST-STING-CT 281-379 in PBS-T containing 3% BSA. The membrane was washed three times with PBS-T and then blotted with HRP-conjugated anti-GST antibody.

**Protein Identification by Mass Spectrometry**

Affinity purified STING complexes were separated on 10% NuPAGE Tris-Glycine gels (Thermo Scientific) and visualized with SimplyBlue SafeStain (Thermo Scientific). Protein bands were excised and de-stained in 50 mM ammonium bicarbonate (ABC) and 50% acetonitrile (ACN) for 10 min. Gel slices were dehydrated in 100% ACN and rehydrated in 50 mM ABC plus 5 mM DTT for 30 min. 50 mM iodoacetamide was added for 60 min. Gel pieces were dehydrated, rehydrated in ABC, and additionally dehydrated before overnight incubation with 12.5 ng/μL trypsin (Promega) in 50 mM ABC at 37 °C. The resulting peptides were extracted in 1% formic acid (FA) at 25°C for 4 hr and then in 0.5% FA/0.5% ACN for 2 hr. After acidification in 1% FA, peptide mixtures were further purified with C18 Zip-tip (Millipore) and analysed by nano–liquid chromatography–mass spectrometry (nLC-MS).

Peptide mixtures were separated on an in-house packed C18 column in silica capillary emitters (resin: 100 Å, 3 μm, MICHROM Bioresources; column: 100 μm ID, 100 mm resin length). A Dionex Ultimate 3000 nanoLC system (Thermo Scientific) provided the LC gradient with 0.1% formic acid as mobile phase A and 0.1% formic acid in acetonitrile as mobile phase B. The following gradient was used: 2% B at 0–15 min, 30% B at 81 min, 35% B at 85 min, 40% B at 87 min, 60% B at 95 min, 80% B at 96–107 min, and 2% B at 108–120 min. Flow rate was 600 nL/min at 0–13.5 min and 250 nL/min at 13.5–120 min.

Peptide eluents were sprayed online with a nano-electrospray ion source (Thermo Scientific) at a spray voltage of 1.5 kV and a capillary temperature of 250°C. High-resolution MS analysis was performed on a QExactive HF-X Quadrupole-Orbitrap Hybrid mass spectrometer (Thermo Scientific), operating in data-dependent mode with dynamic exclusion of 30 s. Full-scan MS was acquired at an m/z range of 300–1650, resolution of 70,000, and automatic gain control target of $3×10^6$ ions. The top 15 most intense ions were subsequently selected for higher-energy collisional dissociation (HCD) fragmentation at a resolution of 17,500, collision energy of 30 eV, and automatic gain control target of $1×10^5$.

Peptide-spectrum matches were performed by the SEQUEST algorithm in Proteome Discoverer (Thermo Scientific), using the *homo sapiens* proteome database (Uniprot, UP000005640) plus common contaminants. Static modification: carbamidomethylation on cysteines; variable modifications: Serine or Threonine phosphorylation, Methionine oxidation, and Glutamine or Asparagine deamination; precursor mass error: 10 ppm; fragment mass error: 0.05 Da; maximum mis-cleavage: 2; peptide false discovery rate: 1%.

**Statistics and reproducibility**

All experiments were independently reproduced at least three times except the STING IP mass spectrometry which was performed once. Strict standards were applied to screen for robust and unbiased results. No statistical methods were used to predetermine the sample size. The investigators were blinded to allocation during imaging and data analysis. Data were presented as mean ± sem. Statistical significance was determined by two-way ANOVA, followed by Turkey's multiple comparison tests.

**Data availability**

The mass spectrometry data have been deposited to the ProteomeXchange Consortium via the PRIDE partner repository under accession number PXD059426. All other data are provided within the paper and its Supplementary Information. Unique materials generated during this study are available from the corresponding author upon request.

#### Supplementary Figure Legends

##### Extended Data Figure 1

**Characterizing STING trafficking dynamics upon cGAMP delivery.**

**(a)** Digitonin-mediated cGAMP delivery triggers robust STING trafficking to perinuclear compartments. BJ cells treated with 100 nM cGAMP were fixed at indicated time points for immunostaining of endogenous STING.

**(b)** *Cis*-Golgi (GM130) appears to wrap around TGN (TGN38) in immunofluorescence. BJ cells treated with 100 nM cGAMP were fixed at indicated time points for co-staining of GM130 and TGN38.

**(c)** GOLGA4 is a more stable TGN marker than TGN38 in cGAMP-stimulated BJ cells. Cells treated with 100 nM cGAMP were fixed at indicated time points for co-staining of GOLGA4 and TGN38.

**(d)** Tracking STING movement through the Golgi using OSBP-PH-GFP as another TGN marker. OSBP-PH-GFP recognizes PI4P on TGN.

**(e)** Brefeldin A disrupts the Golgi complex in BJ cells. BJ cells treated with Brefeldin A (2 μM) for indicated time periods were fixed for co-staining of GM130 and TGN38.

Bar, 10 μm.

**Extended Data Figure 2**

**Mass spectrometry analysis identifies TBK1 and ACBD3 as STING interacting proteins.**

**(a)** Top 500 hits in STING IP mass spectrometry.

**(b)** STING colocalizes with ACBD3 during its trafficking through the Golgi. Bar, 10 μm.

**(c)** Co-immunoprecipitation shows STING interaction with TBK1 and ACBD3 in a cGAMP-dependent manner.

**(d)** Co-immunoprecipitation of endogenous level of Flag-STING with PIKfyve. BJ *STING*$^{-/-}$ cells stably expressing endogenous level of Flag-STING (left) were harvested for IP with anti-Flag-M2 beads (right).

**(e)** PIKfyve shows partial, constitutive colocalization with STING. BJ cells stably expressing Flag-PIKfyve were treated with 100 nM cGAMP and fixed for co-staining of Flag and endogenous STING. Bar, 10 μm.

**Extended Data Figure 3**

**PIKfyve depletion in BJ cells.**

**(a)** BJ cells treated with PIKfyve targeting CRISPR lentivirus develop enlarged endosomes over time. Cells infected with PIKfyve CRISPR lentivirus for 48 hours (Day 0) were subjected to puromycin selection for 24 hours (Day 1). Cells were then checked under microscope and images taken at Day 3, 5 and 7. Cells were seeded for colonization at Day 7. Bar, 10 μm.

**(b)** After 1 month of colonization, most cells with accumulation of enlarged endosomes were lost, but two clones with minor endosome enlargement were obtained. RNAi-mediated depletion of remaining PIKfyve in the two CRISPR clones caused severe accumulation of enlarged endosomes. Bar, 20 μm.

**(c)** Genomic sequencing confirms frame-shift mutations in both PIKfyve alleles from each of the survived sgRNA clones.

**(d)** PIKfyve RNAi did not affect proliferation of BJ PIKfyve CRISPR monoclonal cells within 72 hours. Bar, 20 μm.

**(e)** Complete PIKfyve depletion did not affect Sendai virus (SeV)-stimulated TBK1 phosphorylation in BJ cells. Three days after siRNA transfection, indicated cells were infected with SeV for 3 or 6 hours and whole cell lysates analyzed by immunoblotting.

**(f)** Complete PIKfyve depletion did not affect TNFα-stimulated phosphorylation of IκBα or TBK1. Three days after siRNA transfection, indicated cells were stimulated with TNFα and whole cell lysates harvested at indicated time points were analyzed by immunoblotting.

**(g)** Schematic summary of the strategies used to deplete PIKfyve in BJ cells and their impacts on STING-TBK1 signaling.

**Extended Data Figure 4**

**The PIKfyve inhibitor YM201636 blocks STING-mediated TBK1 activation.**

**(a-b)** Prolonged YM201636 treatment blocks HT-DNA- or cGAMP-stimulated phosphorylation of TBK1 and STING in BJ cells. BJ cells pretreated with DMSO or 5 μM of YM201636 for 18 hours were transfected with HT-DNA (2 μg/ml) for 3 hours (a) or delivered with 100 nM cGAMP for 1-3 hours (b) and whole cell lysates were analyzed by immunoblotting.

**(c-d)** Prolonged YM201636 treatment blocks cGAMP-stimulated TBK1 signaling in L929 (c) and MEF (d) cells. Cells pretreated with DMSO or 5 μM of YM201636 for 18 hours were delivered with 100 nM cGAMP for 1 hour and whole cell lysates were analyzed by immunoblotting.

**(e)** Prolonged YM201636 treatment does not affect SeV-stimulated TBK1 activation in BJ cells. BJ cells pretreated with DMSO or 5 μM of YM201636 for 18 hours were infected with SeV for 6 hours and whole cell lysates were analyzed by immunoblotting.

**Extended Data Figure 5**

**Establishing an *in vitro* liposome-based assay for reconstitution of STING signaling.**

**(a)** The in vitro assay was performed as depicted in Figure 3a. Addition of diC8 PtdIns(3,5)$P_2$ to the in vitro system promotes cGAMP-stimulated phosphorylation of TBK1 and STING, with a substantial amount of activity observed at 25 $\mu$M of the lipid.

**(b,c)** A successful reconstitution of cGAMP-stimulated phosphorylation of TBK1 and STING using cell-derived liposomes requires the addition of cGAMP, diC8 PtdIns(3,5)$P_2$, and ATP (b), as well as expression of STING in COS7 cells (c).

**(d)** A similar assay was also performed using liposomes derived from STING-overexpressing BJ cells.

Compound concentrations in b-d: cGAMP, 1 $\mu$M; diC8 PtdIns(3,5)$P_2$, 25 $\mu$M in solution; ATP, 2 mM.

**Extended Data Figure 6**

**PtdIns(3,5)$P_2$ binding to and regulation of STING.**

**(a)** GST-STING-CT 281-379 does not bind to PIP strips membrane. GST-PLCδ-PH, a positive control that binds to PtdIns(4,5)$P_2$, and GST-STING-CT 281-379 were used in PIP strips assays.

**(b)** Coomassie blue staining of purified 6xHis-EGFP-2xPX.

**(c)** Competition FRET assays performed using 1 μM TMR-PtdIns(3,5)$P_2$, 50 nM STING-mEGFP and increasing concentrations of non-labeled phosphoinositides. Relative binding was calculated as fold changes of FRET-induced loss of EGFP emission.

**(d)** Inhibition of PIKfyve (YM201636) or PI4KB (PI4KIII beta inhibitor 3 and PIK-93) suppresses cGAMP-induced TBK1 phosphorylation. BJ cells were pre-treated with DMSO, YM201636 (4 μM), PI4KIII beta inhibitor 3 (2 μM), or PIK-93 (2 μM) for 18 hours, then stimulated with 100 nM cGAMP for 1 h. Whole cell lysates were harvested for immunoblotting. Note, all inhibitors induced basal LC3 lipidation.

**(e)** Inhibition of PIKfyve or PI4KB suppressed the transcriptional upregulation of *IFNB1* and *CCL5*, an interferon-stimulated gene. BJ cells were pre-treated with DMSO, YM201636 (4 μM), PI4KIII beta inhibitor 3 (2 μM), or PIK-93 (2 μM) for 18 hours, stimulated with 100 nM cGAMP for 2 h, and followed by RNA extraction and qRT-PCR. Mean ± sd; n = 5. Two-way ANNOVA, followed by Turkey's multiple comparisons tests.

**Extended Data Figure 7**

**PtdIns(3,5)$P_2$-binding supports both the canonical and noncanonical functions of STING**

**(a)** Sequence alignment showing basic residues in the N-terminus and the cytosolic loop of STING.

**(b)** Data summary of STING induced self-quenching of TMR-PtdIns(3,5)$P_2$. Emission of TMR-PtdIns(3,5)$P_2$ at 574 nm and 590 nm was captured in the absence or presence of 100 nM purified STING without mEGFP tag. Mean ± SEM; n = 3.

**(c)** FRET between 250 nM TMR-PtdIns(3,5)$P_2$ and 500 nM STING-mEGFP or indicated mutants.

**(a)** The PtdIns(3,5)$P_2$-binding mutant of STING (K20A/R71A) does not respond to PtdIns(3,5)$P_2$ in in vitro reconstitution of STING-TBK1 signaling using COS7-derived liposomes. cGAMP: 1 μM; diC8 PtdIns(3,5)$P_2$: 25 μM in solution; ATP: 2 mM.

**(d)** The PtdIns(3,5)$P_2$-binding mutant of STING shows defects in inducing TFEB activation and LC3 lipidation. U2OS cells expressing STING or the K20A/R71A mutant were treated with 100 nM cGAMP for 2 h, and whole cell lysates were harvested for immunoblotting.

**(e)** Quantification of LC3-II intensities normalized to GAPDH. Mean ± sd; n = 3. Two-way ANNOVA, followed by Turkey's multiple comparisons tests.

**Extended Data Figure 8**

**The PIKfyve inhibitor YM201636, which incompletely blocks PIKfyve activity, suppresses STING signaling activation without affecting STING trafficking from the ER.**

**(a)** Morphological changes of BJ cells after indicated treatments. YM201636 was used at 5 μM. siRNA was transfected three days before imaging.

**(b)** YM201636 treatment does not block STING trafficking to perinuclear endosomes. BJ cells pretreated with DMSO or YM201636 (5 μM) for 18 hours were stimulated with 100 nM cGAMP and then fixed at indicated time points for co-staining of endogenous STING and TBK1. Bar, 10 μm. Note that the imaging only shows the recruitment of TBK1 but cannot tell how "stable" the interaction is. YM201636 strongly reduced the STING-TBK1 co-IP (Extended Data Fig. 9b), suggesting that the observed TBK1 recruitment in immunofluorescence are based on less stable, weak interactions.

**(c)** Top: Schematic illustration of the differences between two mEGFP fusion proteins of STING. In STING-110mEGFP, the sequence of mEGFP was inserted to the second luminal loop of STING after amino acid 110. The structure of the fusion proteins is predicted by AlphaFold. Bottom: STING-110mEGFP, but not STING-mEGFP, rescues cGAMP-stimulated TBK1 phosphorylation in BJ STING-knockout cells. BJ wild type and STING knockout cells reconstituted or not with STING-mEGFP or STING-110mEGFP were stimulated with cGAMP and then analyzed by western blot.

**(d)** YM201636 blocks STING-110mEGFP-mediated TBK1 activation without affecting its trafficking from the ER to perinuclear compartments. BJ STING knockout cells reconstituted with STING-110mEGFP were pretreated with DMSO or YM201636 (5 μM) for 18 hours and then stimulated with or without 100 nM cGAMP. Fluorescence images were taken 60 min after cGAMP exposure, followed by lysis of the same cells for immunoblotting using indicated antibodies. Bar, 20 μm.

**(e)** YM201636 suppresses STING-dependent TBK1 phosphorylation shown by immunofluorescence images. BJ STING-KO cells stably expressing STING-110mEGFP were pre-treated with DMSO or 4 μM YM201636 (18 h), stimulated with 100 nM cGAMP for 1 h. Cells were then fixed for the staining of endogenous p-TBK1. DAPI stains the nuclei. Bar, 10 μm.

**(f)** Quantification of p-TBK1 intensities in (e) above threshold (A.U.). Mean ± sd; n = 3. Two-way ANNOVA, followed by Turkey's multiple comparisons tests.

**Extended Data Figure 9**

**STING-stimulated TBK1 phosphorylation requires TBK1 kinase activity and PtdIns(3,5)$P_2$.**

**(b)** TBK1 K38A mutant reconstituted in BJ TBK1 knockout cells cannot be phosphorylated in response to cGAMP or HT-DNA treatment. BJ wild type and TBK1/IKKε double knockout cells reconstituted or not with wild type or K38A TBK1 were stimulated with 100 nM cGAMP for 1 hour or transfected with 2 μg/ml HT-DNA or lipofection alone for 3 hours. Whole cell lysates were harvested for immunoblotting analysis. Stable re-expression of wild-type (WT) or K38A TBK1-Flag in TBK1/IKKε double knockout cells was achieved by lentiviral infection.

**(c)** YM201636 treatment inhibits STING/TBK1 interaction in co-IP. BJ cells stably expressing Flag-STING were pretreated with DMSO or 5 μM YM201636 for 18 hours and stimulated with cGAMP for 1 hour, followed by IP of whole cell lysates with anti-Flag M2 affinity gel. The cell lysate and IP samples were analyzed by immunoblotting.

**(d)** After a liposome-based reaction as depicted in Figure 3a, most TBK1, p-TBK1 and STING were associated with the membrane fraction. After 10 min of reaction at 37 °C, half of sample was saved as total input and the other half was centrifuged at 20,000 *xg* for 10 min at 4 °C. Pellet and supernatant were diluted with SDS loading buffer to the same volume of the total sample and analyzed by immunoblotting. cGAMP: 1 μM; diC8 PtdIns(3,5)$P_2$: 25 μM in solution; ATP: 2 mM.

**Extended Data Figure 10**

**Summary of PtdIns(3,5)$P_2$-mediated STING trafficking and signaling activation.**

Summary of experimental evidence supporting a key role for PtdIns(3,5)$P_2$ in STING trafficking and TBK1 activation.

**Table S1. Primer, siRNA, and gRNA sequences for human STING and TBK1**

| Oligonucleotide name | Sequence |
|---|---|
| STING R281Q | CAATACAGTCAAGCTGGCTTTAGCCAGGAGGATAGGCTTGAGCAGGCC |
| STING K289Q | GGAGGATAGGCTTGAGCAGGCCCAACTCTTCTGCCGGACACTTGAGGAC |
| STING R293Q | GCTTGAGCAGGCCAAACTCTTCTGCCAGACACTTGAGGACATCCTGGCAG |
| STING K289Q/R293Q | GGAGGATAGGCTTGAGCAGGCCCAACTCTTCTGCCAGACACTTGAGGACATCCTGGCAG |
| STING R310Q | CCCTGAGTCTCAGAACAACTGCCAACTCATTGCCTACCAGGAACCTGC |
| STING R331Q | CTCGCTGTCCCAGGAGGTTCTCCAGCACCTGCGGCAGGAGGAAAAGG |
| EGFP A206K | GACAACCACTACCTGAGCACCCAGTCCAAACTGAGCAAAGACCCCAACGAGAAGCG |
| STING R76Q | GGCTGAGGAGCTGCGCCACATCCACTCCCAGTACCGGGGCAGCTACTGGAGGACTGTG |
| STING R76,78Q | GCTGAGGAGCTGCGCCACATCCACTCCCAGTACCAGGGCAGCTACTGGAGGACTGTGCGG |
| STING R83,86Q | TCCAGGTACCGGGGCAGCTACTGGCAGACTGTGCAGGCCTGCCTGGGCTGCCCCCTCCG |
| STING-mEGFP R375A | GTGGAATGGAAAAGCCCCTCCCTCTCGCCACGGATTTCTCTCTCGAGATGGTGAGC |
| STING 71Q72,74L | GTCTGCAGCCTGGCTGAGGAGCTGCAGCTGATCCTGTCCAGGTACCGGGGCAGCTACTGG |
| STING H72L/R76Q | CAGCCTGGCTGAGGAGCTGCGCCTGATCCACTCCCAGTACCGGGGCAGCTACTGGCAGAC |
| STING R71A_F | CACATCCACTCCAGGTACCGG |
| STING R71A_R | CCTGGAGTGGATGTGGGCCAGCTCCTCAGCCAGGCTG |
| STING K20A_F | ATGCCCCACTCCAGCCTGCATCCATCCATCCCGTGTCCCAGGGGTCACGGGGCCCAGGCGGCAGCCT TGGTTCTGCTGA |
| STQ273AA277Q_F | GCATACAGTCAACAAGGCTTTAGCCGGGAGGATAGG |
| STQ273AA277Q_R | TTGTTGACTGTATGCTGACATGGCAAACAAAGTCTGCAAG |
| STING sgRNA resistant mutations | GCTTGCCCTCCTGGGCCTCTCACAAGCGCTGAACATCCTCCTGGGCCTC |
| TBK1 sgRNA resistant mutations | TATACACTGTTTTAGAAGAACCGAGCAACGCTTATGGACTACCAGAATCTGAA |
| TBK1 sgRNA target | GAAGAACCTTCTAATGCCTA |
| IKKε sgRNA target | TTTCAGGGCGTGTTGGGCGC |
| PIKfyve sgRNA target | AGACCGACTGCAGTTGCTGA |
| Control siRNA | AGGUAGUGUAAUCGCCUUG |
| PIKfyve siRNA #1 | CCAACUGCAUUGUAGGAAA |
| PIKfyve siRNA#2 | GCUAGUGAGCACUAGAAUU |

### Extended Data Fig. 1

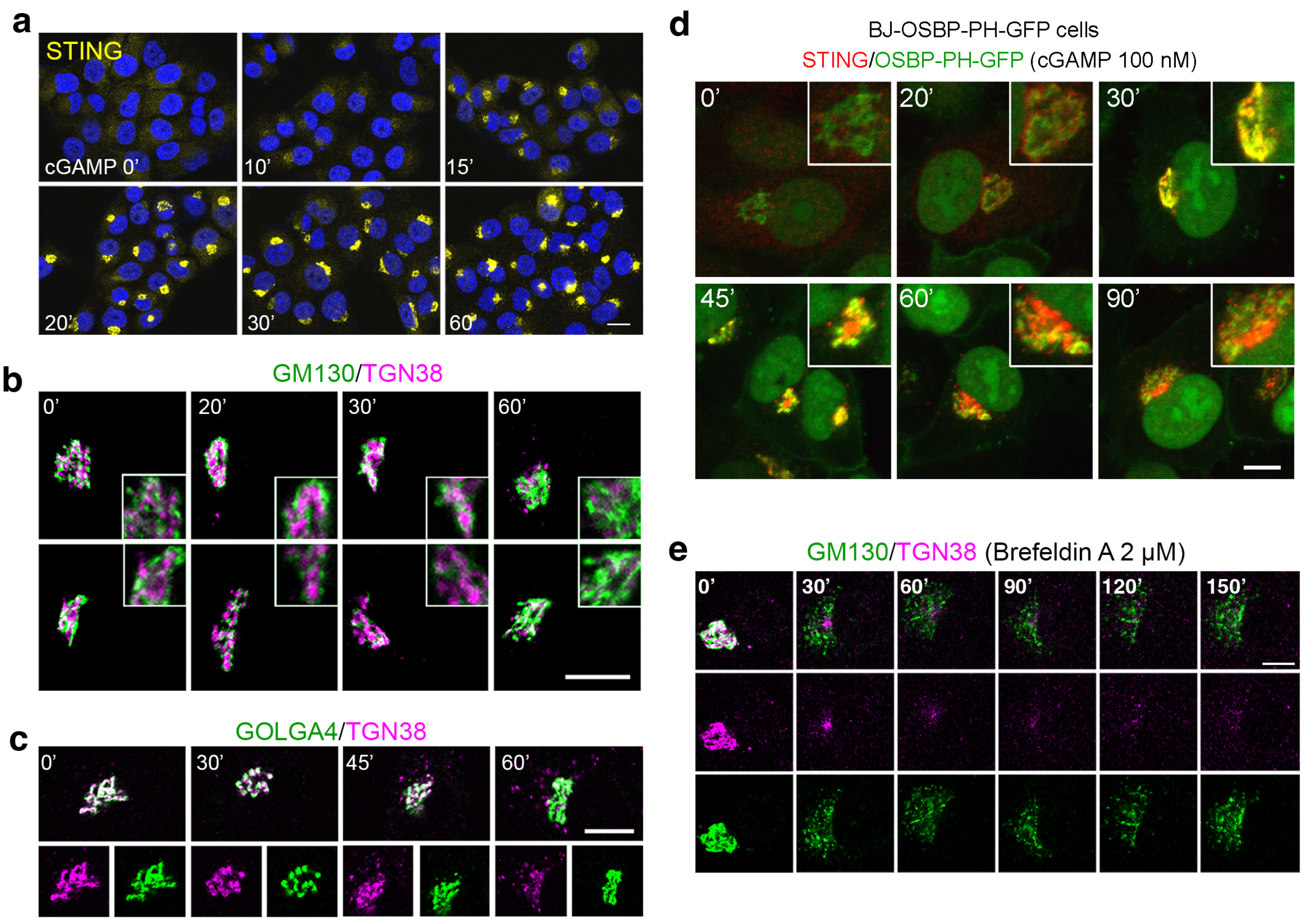

#### Extended Data Fig. 2

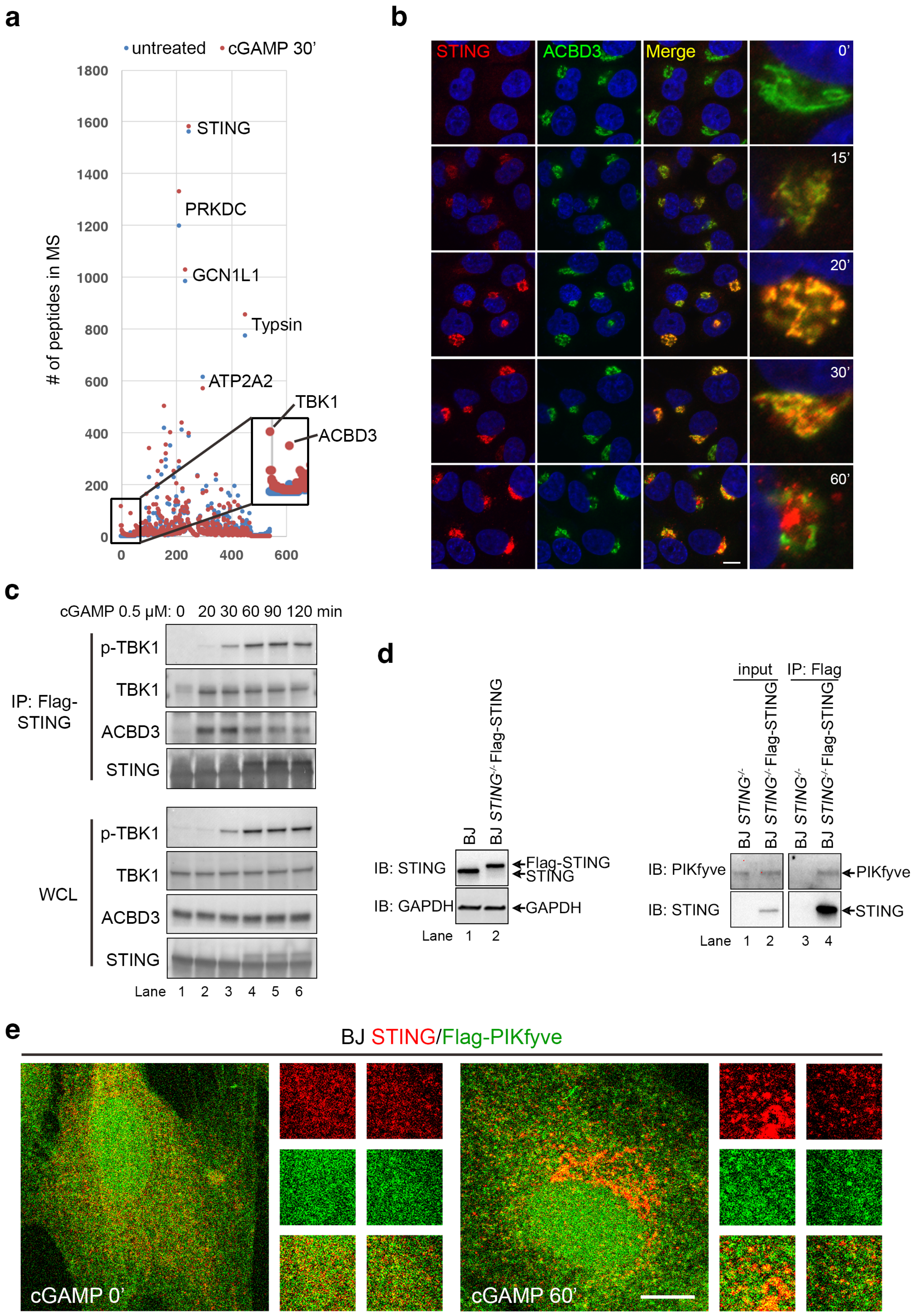

### Extended Data Fig 3

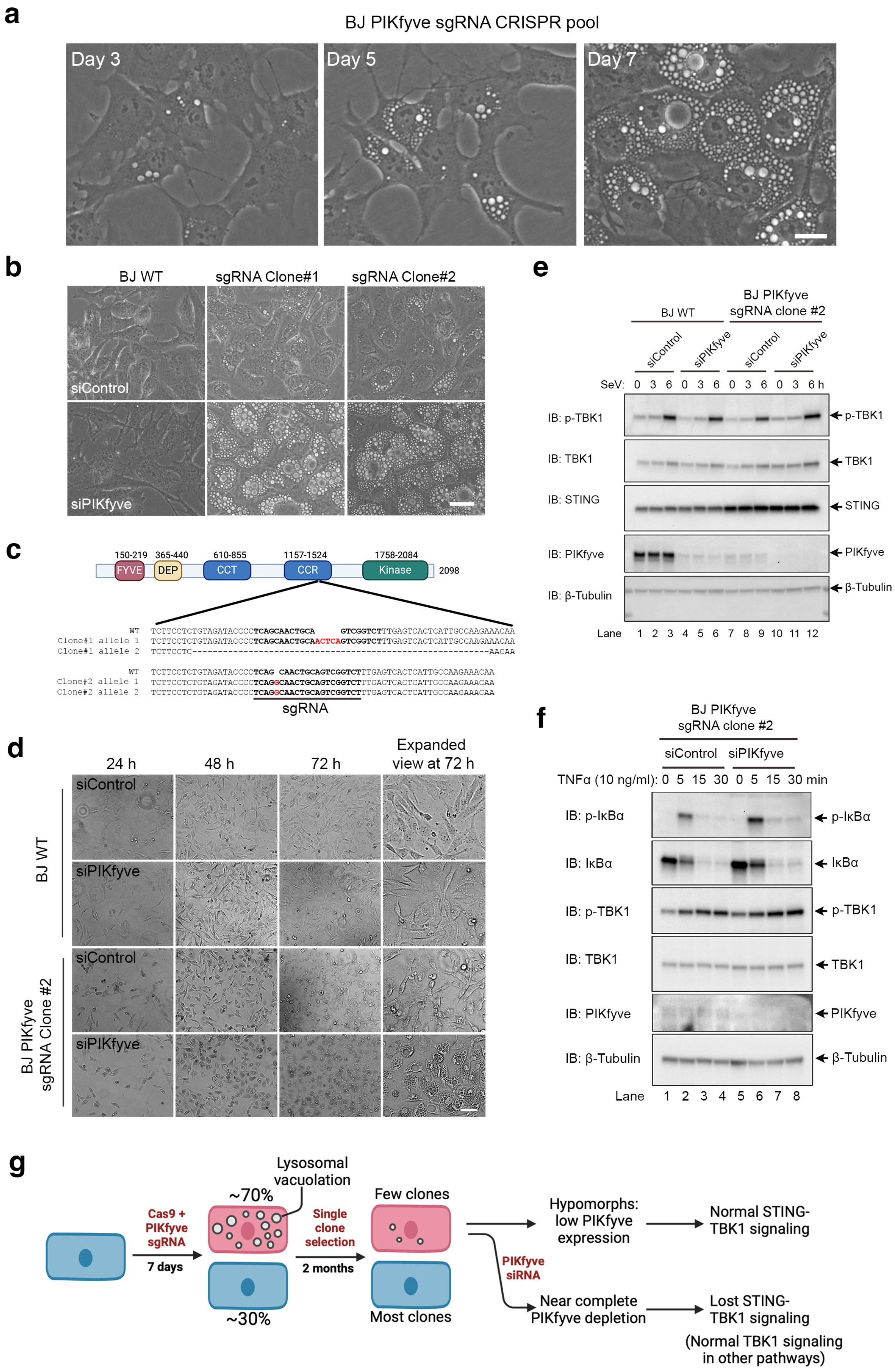

### Extended Data Fig 4

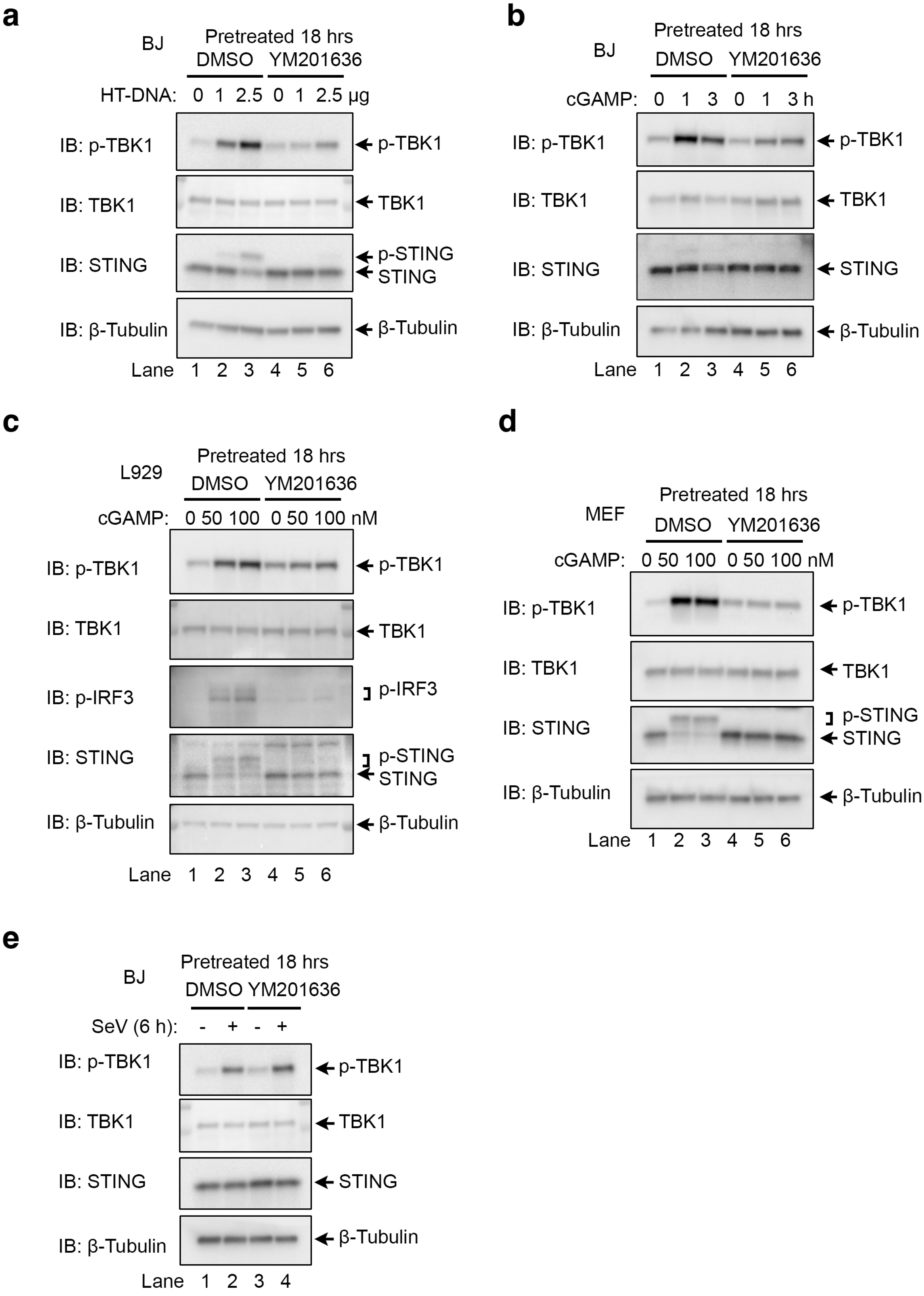

#### Extended Data Fig. 5

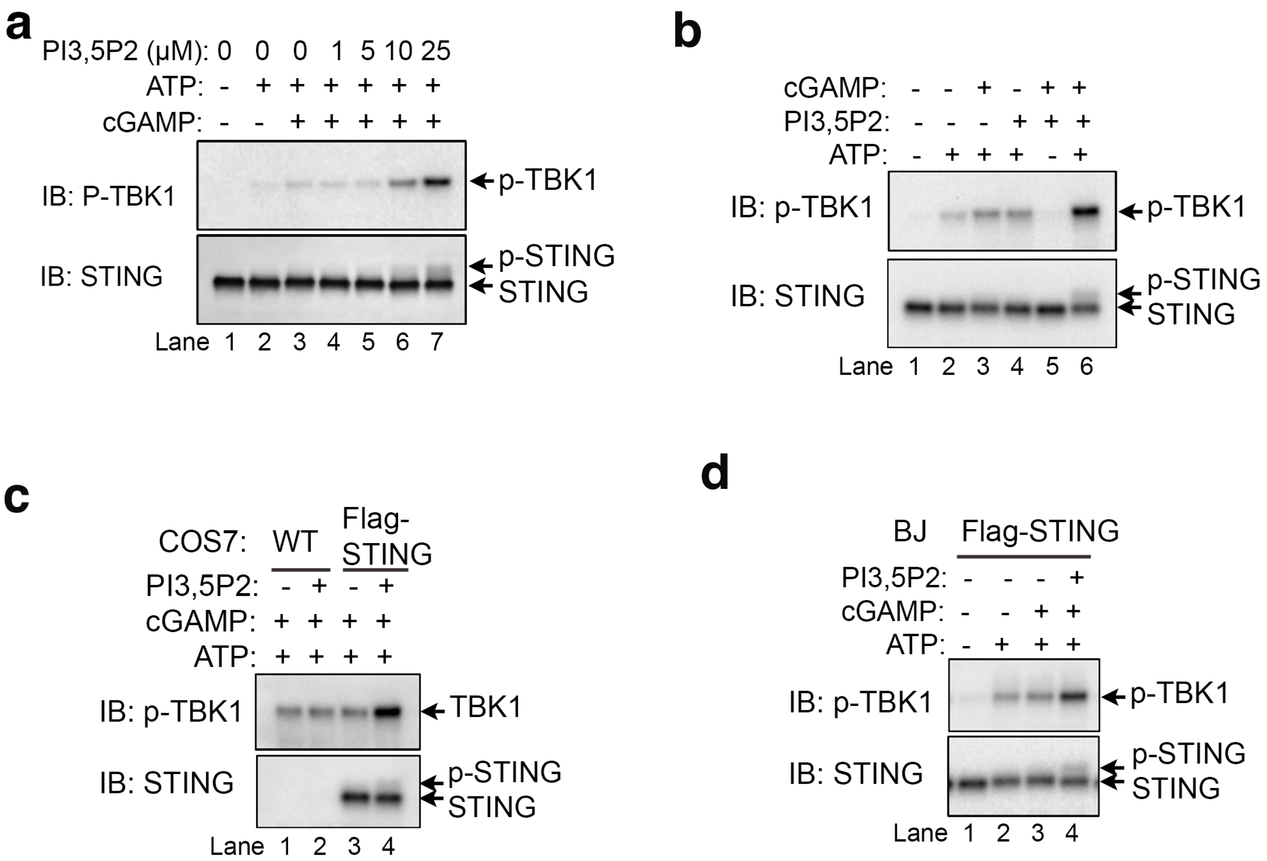

**Extended Data Fig. 6**

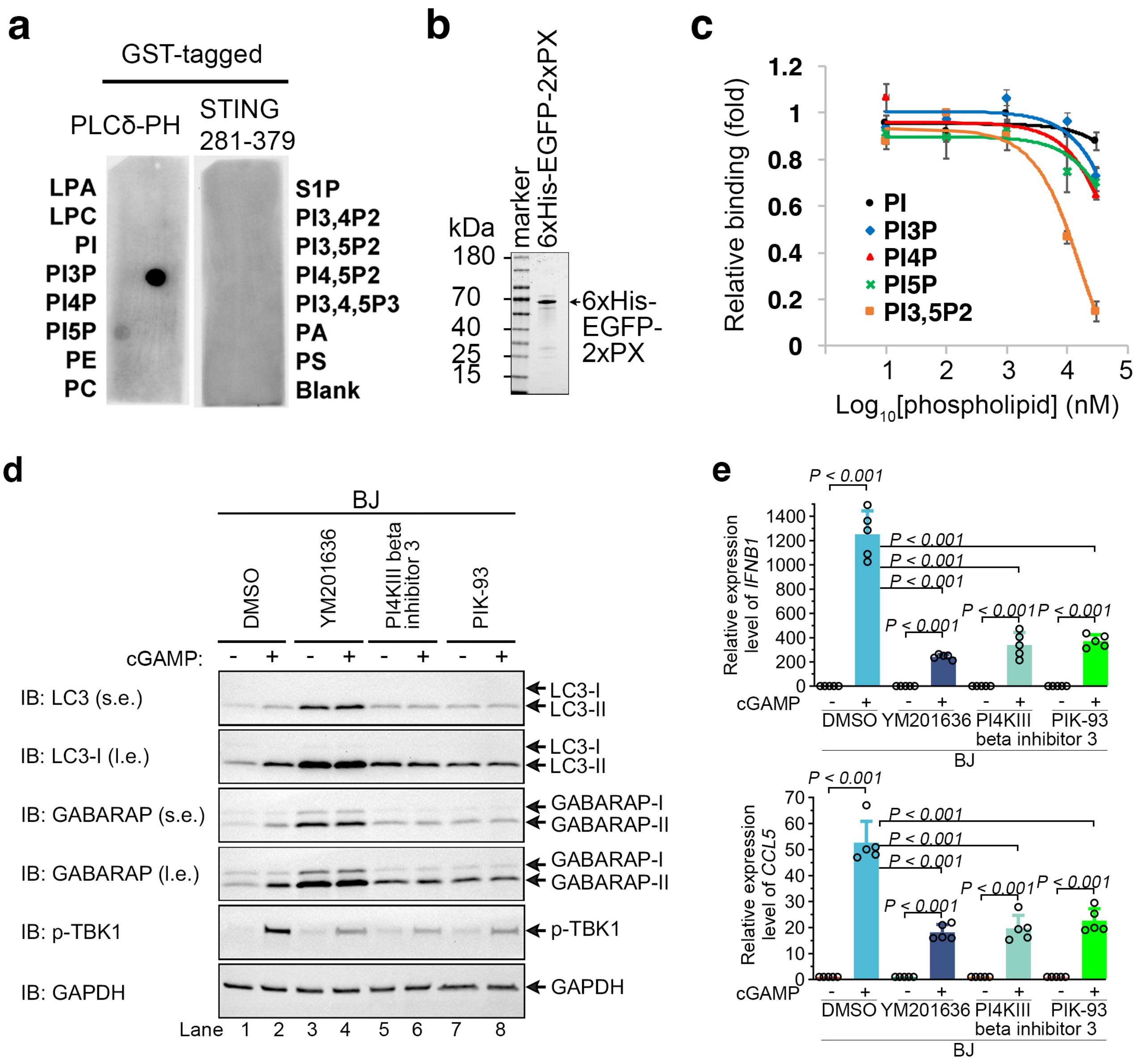

### Extended Data Fig 7

**a**

```
                             N-terminus        Loop pocket
Conservation index:          5 99 596 6        9 95995 9 5599 959956 595
                                  14 16   20              71  74 76 78    83  86
Homo_sapiens                 PSIPCPRGHGAQKA    LNGVCSLAEELRHIHSRYRGSYWRTVRACLG
Pan_troglodytes              PSIPCPRGHGAQKA    LNGVCSLAEELRHIHSRYWGSYWRTVRACLG
Canis_lupus_familiaris       PSIPRPRGTRAQKA    FKGVCYLTEELCHLHSRYQGSYWRATRACLG
Bos_taurus                   PSIPQPRGLRAQKA    IKGICSLAEELCHVHSRYHGSYWRAVRACLC
Mus_musculus                 PAIPRPRGHRSKYV    LKNLCCLAEELCHVQSRYQGSYWKAVRACLG
Rattus_norvegicus            PSIPRPRSYRFKLA    LKKLCCLAEELCHVQSRYQGSYWKAVRACVG
Xenopus_tropicalis           QIIPERRGKRATKM    VTCACELTEEIKHLHTRYNGHYWRALKASFN
Consensus:                         R                      H    R         +  +
```

**b**

STING induced TMR-PI3,5P2 quenching

| n = 5 | STING | 0 nM | 100 nM |
|---|---|---|---|
| 574 nm | TMR Emission | 100% | 96.3% ± 0.9% |
| | TMR quenching | 0 | 3.72% ± 0.9% |
| 590 nm | TMR Emission % | 100% | 95.6% ± 1.2% |
| | TMR quenching % | 0 | 4.43% ± 1.2% |

**c**

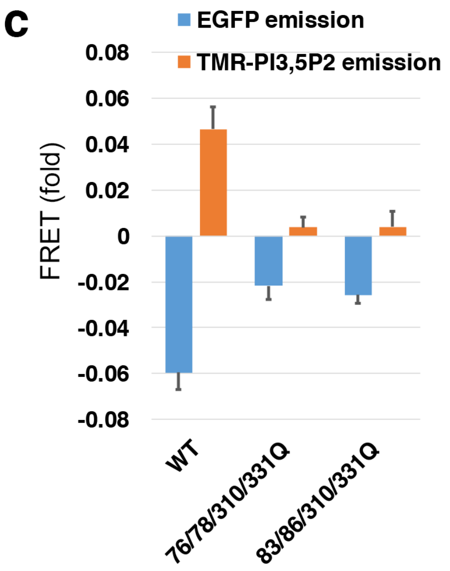


**d**

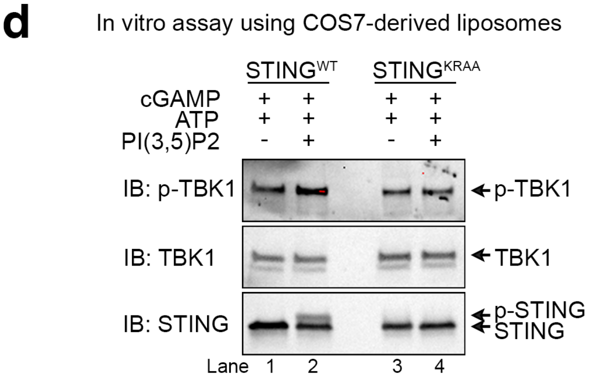


**e**

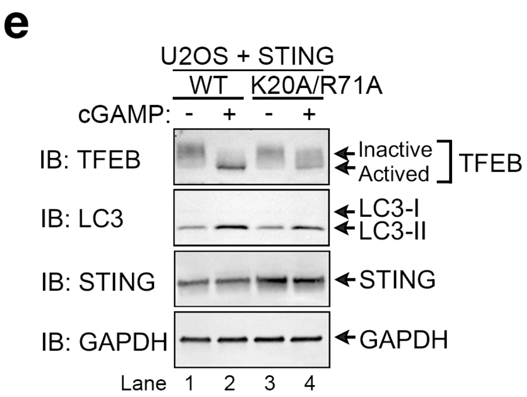


**f**

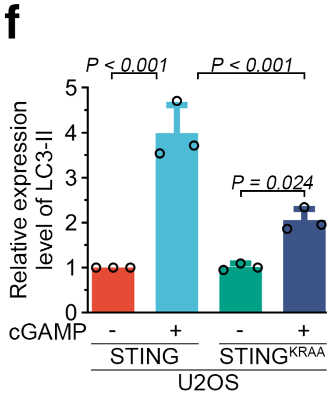

#### Extended Data Fig 8

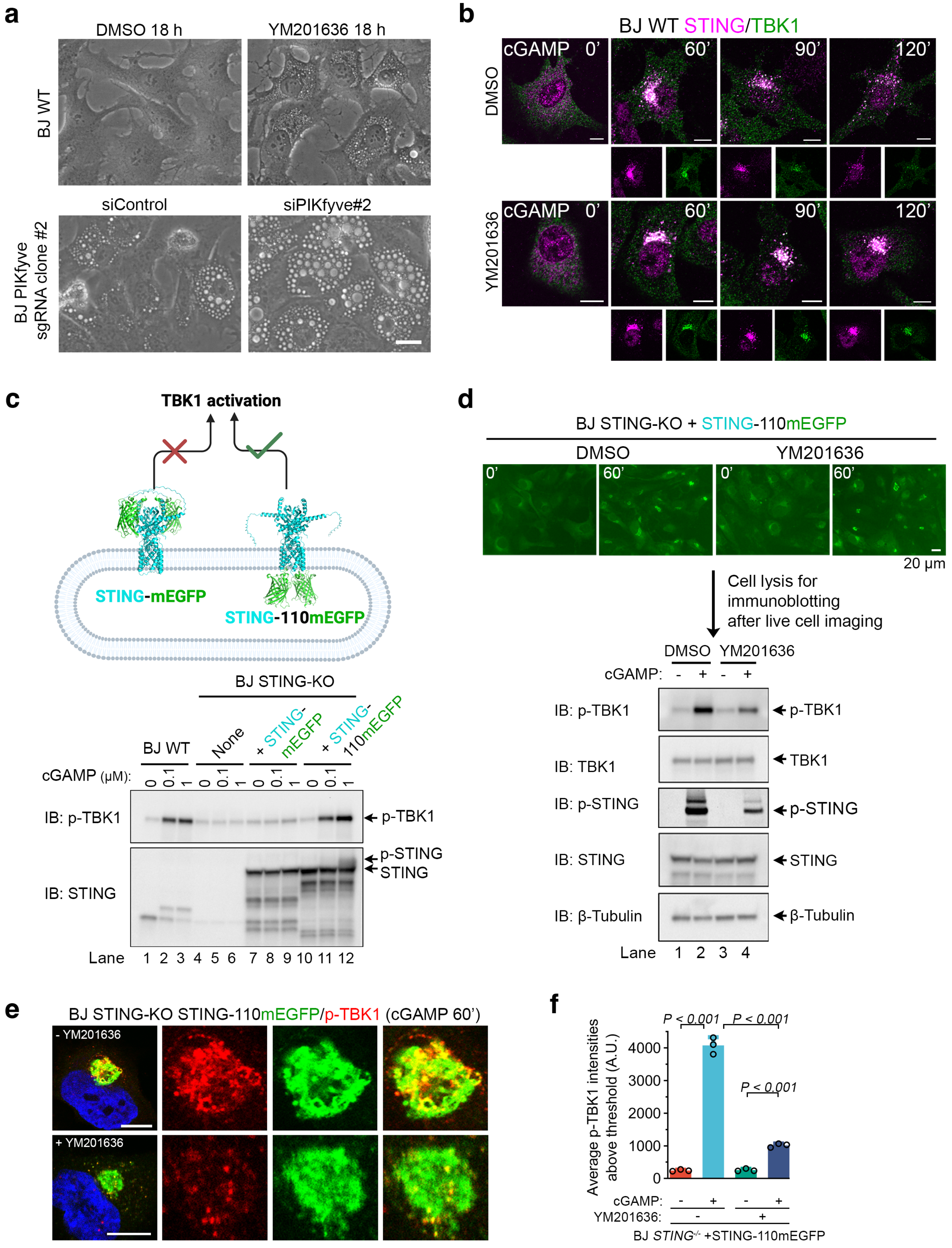

### Extended Data Fig 9

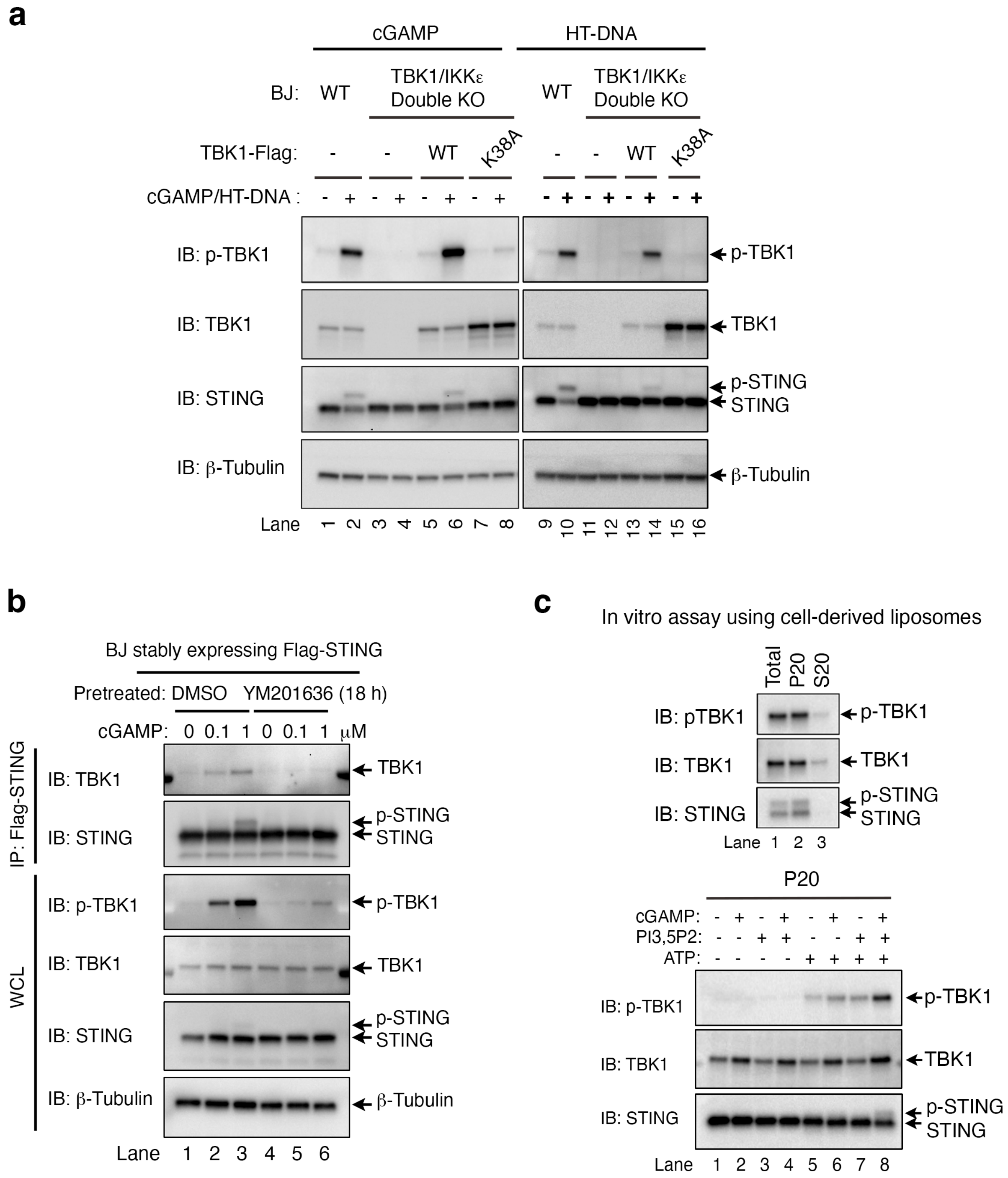

#### Extended Data Fig 10

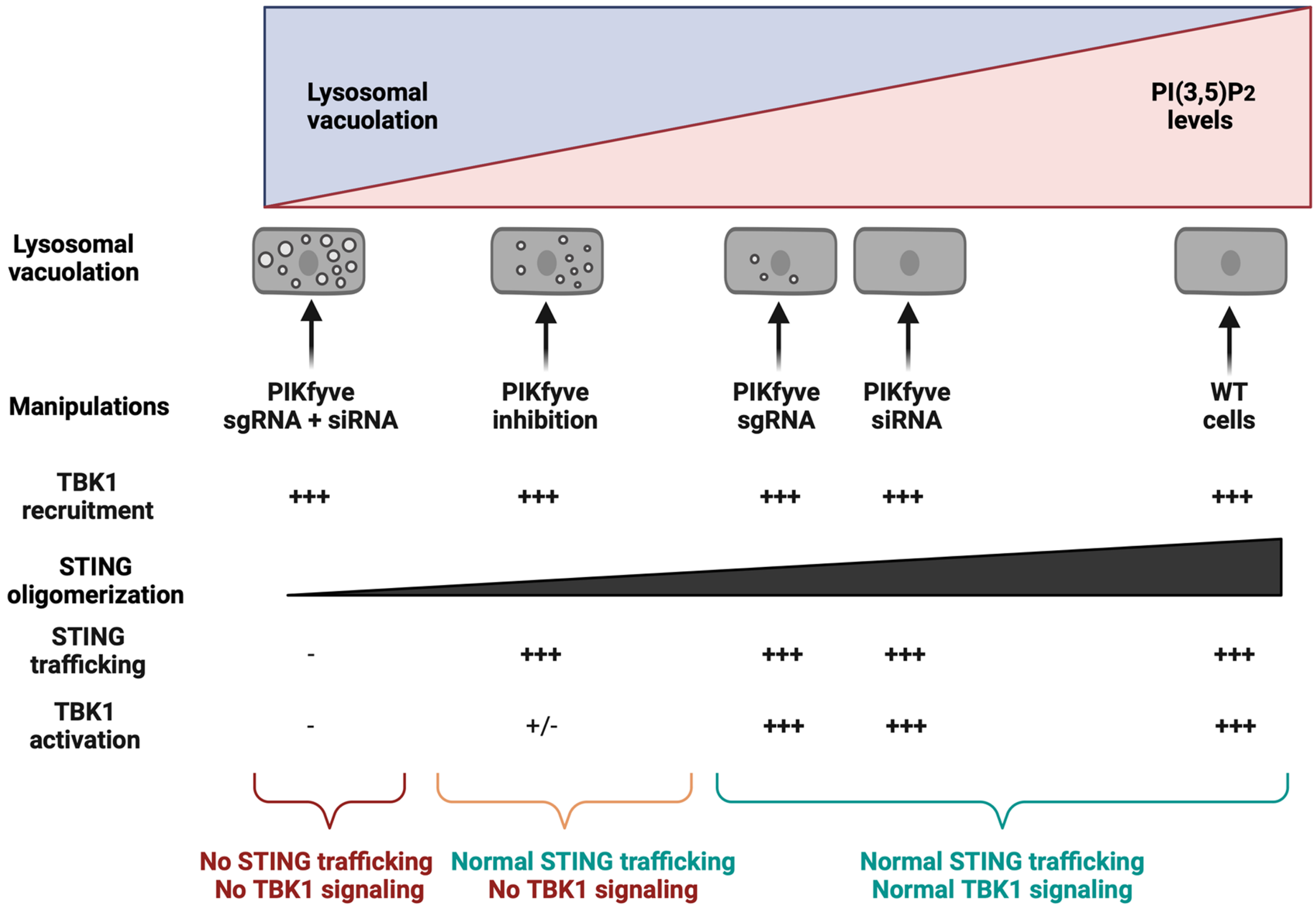
